# Marine heatwaves drive range contraction and alternative states of kelp forests at their warm limit

**DOI:** 10.1101/2025.10.17.682914

**Authors:** Nur Arafeh-Dalmau, David S. Schoeman, Gabriela Montaño-Moctezuma, Guillermo Torres-Moye, Kyle C. Cavanaugh, Adrian Munguía-Vega, Octavio Aburto-Oropeza, Jessica A. García-Pantoja, Carolina Olguín-Jacobson, Fiorenza Micheli

## Abstract

Marine heatwaves are transforming ecosystems, yet their role in driving alternative states—and the conditions that enable these transitions—remains poorly understood. Using 30 years of satellite and underwater data, we assessed the impact of the 2014–2016 Pacific marine heatwaves on giant kelp forests (*Macrocystis pyrifera*) at their warm range limit in Mexico. By 2016, 88% of forests were lost, with limited recovery by 2023, including an 80 km range contraction at the southern edge. Surveys revealed three alternative states: replacement by heat-tolerant palm kelp (*Eisenia arborea*) in warmer regions; urchin barrens due to predator overfishing; and, unexpectedly, persistent giant kelp near the southern limit where high temperatures coincide with low human pressure. Pre-existing conditions, such as high urchin and palm kelp densities, shaped these outcomes. These findings show that responses to marine heatwaves are shaped by local ecological and human contexts, requiring tailored climate-adaptation strategies to promote resilience.

## Introduction

Climate change is reshaping marine ecosystems at all levels of biological organization, from gene expression to species interactions and ecosystem structure and function (Cooley et al., 2022; Poloczanska et al., 2013; Poloczanska et al., 2016). Rising ocean temperatures are driving shifts in species distributions (Pecl et al., 2017), with many species moving poleward, displacing temperate species as they track suitable habitats (Vergés et al., 2014). Alongside these gradual warming trends, extreme events like marine heatwaves (MHWs) are causing mass die-offs and range shifts in marine species and ecosystems (Garrabou et al., 2022; Ling & Keane, 2024; McPherson et al., 2021; Sanford et al., 2019; Smith et al., 2023; Wernberg et al., 2016). Despite these well-documented impacts, less attention has been given to whether extreme events may drive ecosystem shifts to alternative stable states and the mechanisms that drive these shifts. This is a critical knowledge gap, as alternative states differ profoundly in biodiversity, ecosystem functioning, and services, and recovery may be slow or absent beyond ecological tipping points (Armstrong McKay et al., 2022; Graham, 2004; Wernberg et al., 2025). Thus, understanding how extreme events reshape ecosystems is crucial to developing effective climate adaptation strategies (Arafeh-Dalmau et al., 2023).

The frequency and intensity of large-scale, prolonged MHWs have increased (Oliver et al., 2018), leading to widespread losses of habitat-forming species like corals, kelps, mangroves and seagrasses (Arafeh-Dalmau et al., 2019; McPherson et al., 2021; Smale, 2020; Smale et al., 2019; Starko et al., 2025; Wernberg et al., 2016; Wernberg et al., 2025). This loss is driving dramatic ecosystem shifts, often resulting in regime changes and alternative stable states that persist (McPherson et al., 2021; Wernberg, 2021) and undermine the ecosystem services provided by biodiversity (Smith et al., 2021; Villaseñor-Derbez, 2024). Kelp forests, which cover over 30% of the world’s coastlines (Jayathilake & Costello, 2021), are particularly vulnerable (Arafeh-Dalmau et al., 2020; Smith et al., 2023), since MHWs can reduce kelp biomass, causing sea urchins to shift from being passive detritus feeders to active kelp grazers, resulting in long-lasting shifts between kelp-dominated and urchin-dominated states (McPherson et al., 2021). The ability of ecosystems to recover from heatwaves is influenced by pre-existing conditions and ecological mechanisms, such as species competition (Wernberg et al., 2019) and predator-prey imbalances (Kumagai et al., 2024; Ling et al., 2009), which can either reinforce or resist transitions to alternative states (Dudgeon et al., 2010; Filbee-Dexter & Scheibling, 2014; Ling et al., 2015). Human-induced stressors, such as overfishing, further erode the resilience of marine ecosystems and exacerbate the impacts of climate change (Baum et al., 2023; Donovan et al., 2021; Gove et al., 2023; Kumagai et al., 2024; Ling & Keane, 2024). While research on how MHWs impact habitat-forming species is expanding (K. E. Smith et al., 2024), a significant knowledge gap exists in understanding the conditions that lead to alternative stable states and whether these shifts are reversible after such events.

Collecting data in marine environments is challenging, often relying on *in situ* underwater observations that limit our understanding of long-term, large-scale responses to extreme events. For floating kelp forests and other habitats visible in aerial imagery, remote sensing can bridge this gap by providing decades of large-scale data on their distribution and dynamics (Bell et al., 2023; Cavanaugh et al., 2021). Because remote sensing cannot capture the ecological mechanisms that drive ecosystem changes, these technologies must be integrated with long-term field observations of species and their interactions—such as predation, herbivory, facilitation and competition—that shape climate-driven regime shifts and ecosystem trajectories (Pinsky et al., 2020).

Here, we used over 30 years of quarterly remote-sensing data (Bell et al., 2023) to track the impact and recovery of giant kelp (*Macrocystis pyrifera)* forests along 1,605 km of coastline in Baja California, Mexico from a series of MHWs in 2014–2016. We then assessed the post-heatwave status of kelp forests and the possible drivers of alternative states through extensive field expeditions in 2022 and 2023, each sampling 32–35 reefs spanning ∼800 km of a temperate– subtropical transition zone. These recent data were supplemented with historical field surveys dating back to 1993 to determine whether pre-existing conditions explained the observed changes. We found that Mexico has lost 50% of its giant kelp forests, including an 80 km range contraction at the species’ southern range limit. MHWs shifted forests into three alternative states dominated by sea urchins, giant kelp, or the heat-tolerant palm kelp *Eisenia arborea*, with their emergence shaped by pre-existing conditions. Contrary to expectations, the most persistent forests occurred near the southern edge, where high predator and low urchin biomass prevailed, highlighting that effective fisheries management and ecosystem protection enhance resilience to climate extremes.

## Methods

The study area comprises the distribution of giant-kelp forests along the Pacific coastline of Mexico (Figure 1). The region extends from the Mexico-USA international border in the north (∼32.53° N), Baja California, to the southern distribution limit of giant kelp in the northern hemisphere in Punta Prieta (∼27° N), Baja California Sur, Mexico. This region represents the southern tip of the California Current System, one of Earth’s most productive ecosystems (Checkley Jr & Barth, 2009). The marine resources and ecosystems in this region were impacted during 2014–2016 by the strongest and longest MHWs ever detected in the satellite record (Arafeh-Dalmau et al., 2019; Beas-Luna et al., 2020; Cavanaugh et al., 2019; Micheli et al., 2024; Villaseñor-Derbez, 2024). We divided the study region into two sub-ecoregions: northern Baja California (32.53–29.73° N), which is characterized by relatively colder waters and assemblages similar to southern California, and central Baja California (29.72–27° N), which has warmer average SST and contains the distribution limit of many species (Arafeh-Dalmau et al., 2023; Lonhart et al., 2019; Ramirez-Valdez et al., 2015). In Baja California, giant kelp co-exist with other subcanopy species, such as the warmer-affinity *Eisenia arborea* that extends further south than *M. pyrifera* (Edwards & Hernandez-Carmona, 2005). Three main herbivore species that consume kelp species are present in the region: *M. franciscanus*, *S. purpuratus*, and *C. coronatus*. These taxa co-exist throughout the study region, although *S. purpuratus* and *M. franciscanus* have a cooler affinity than *C. coronatus*. In both regions, the primary predators of sea urchins are California sheephead (*S. pulcher*) and spiny lobster (*P. interruptus*).

**Figure 1.**
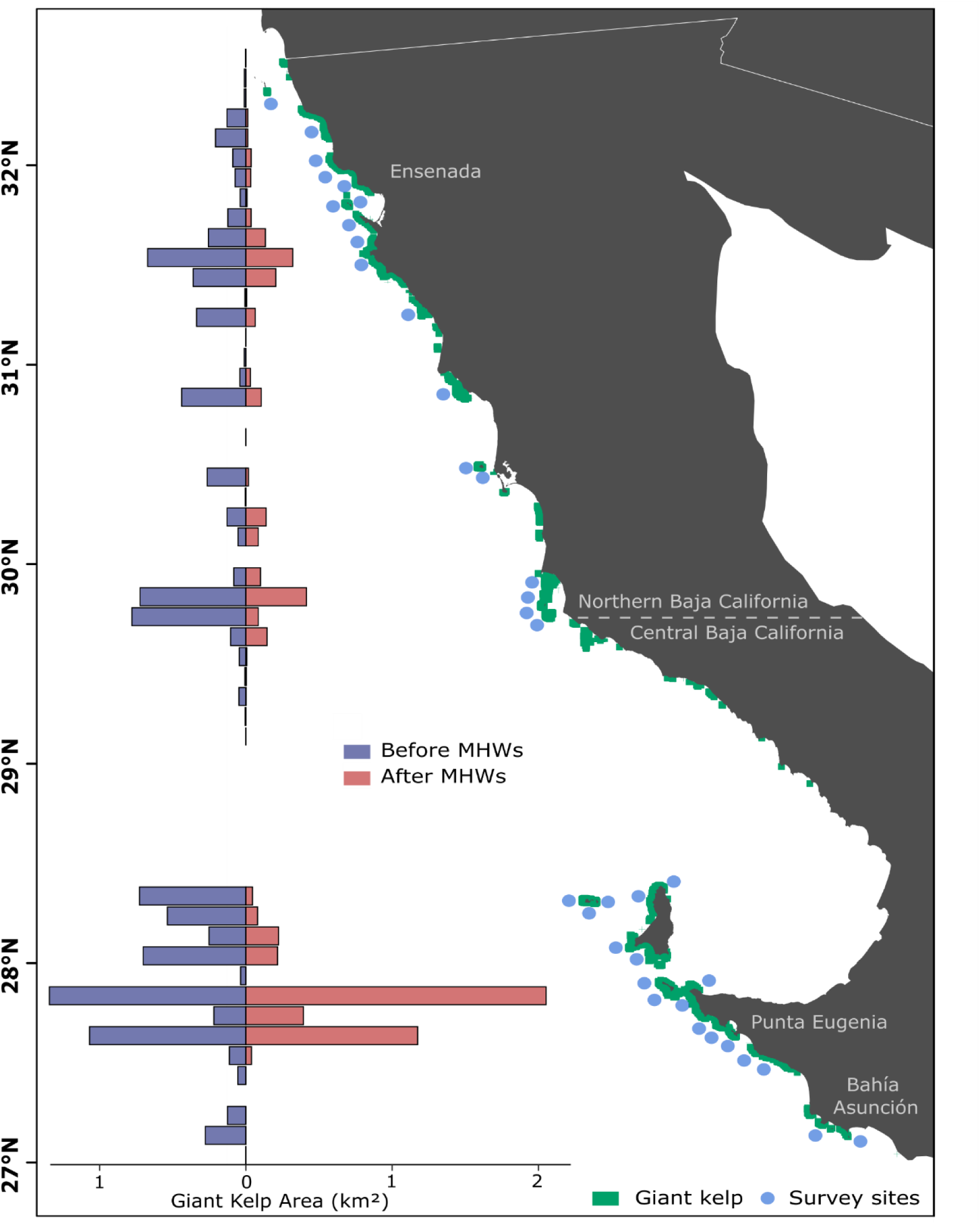
- Kelp forests in Baja California before and after the 2014–2016 MHWs. Bar plots represent the mean annual kelp area (km^2^) from remote-sensing estimates before (2000–2013) and after (2017–2023) MHWs, binned at 0.1°. Blue dots represent underwater survey sites in 2022 and 2023, plotted slightly offshore of their true locations to avoid obscuring kelp distribution. Green polygons represent the distribution of giant kelp forests in Mexico.

### Giant kelp remote-sensing time-series

We used a timeseries developed from 40 years of satellite-based estimates of kelp area at a 30-m-grid resolution to map kelp forests and analyze dynamics in the Peninsula of Baja California, Mexico. This published dataset provides quarterly estimates of kelp canopy area from 1984 to present (Bell et al., 2023) and can be visualized on https://kelpwatch.org. The dataset estimates kelp canopy from three Landsat sensors: Landsat 5 Thematic Mapper (1984–2011), Landsat 7 Enhanced Thematic Mapper+ (1999–present), and Landsat 8 Operational Land Imager (2013– present).

We treated the remote-sensed time series differently for each analysis. First, we aggregated the dataset at 0.1° intervals of latitude to visualize and assess latitudinal changes in mean kelp area (km^2^) before (2000–2013) and after (2017–2023) the MHWs.

Next, we aggregated the dataset into 1-km^2^ pixels to estimate the mean annual kelp area per pixel for each year from 2000–2023 in the central (n = 416 pixels) and northern Baja California (n = 470 pixels) ecoregions, to assess whether kelp forests were impacted by and recovered from the 2014–2016 MHWs, and whether the response varied across ecoregions. For this analysis, we classified each year as either before (2000–2013), during (2014–2016), or after (2017–2023) the MHWs.

Finally, we selected three areas that today are dominated by alternative kelp-forest stable states (i.e., giant kelp forests, palm kelp forests, urchin barrens) and aggregated the data into 0.5° latitudinal bins: Ensenada (31.75–32.25° N); Punta Eugenia (27.5–28° N); and Bahía Asunción (27–27.5° N). For these three regions, we estimated the percentage change in annual kelp coverage over the past 10 years (2014–2023) compared to the baseline mean coverage (2000–2013) before the MHWs. We also developed a time series for each of the three regions for each quarter of the year for the entire study period to assess whether preconditions in these regions determined the subsequent response of kelp forests to the MHWs. While 0.5° bins were used to standardize regional comparisons and assess temporal trends, for Ensenada we used a broader area (31.63– 32.53° N) for the percentage change analysis to capture the full extent of kelp loss during and after the MHWs in the region.

For all data treatment, we followed previous approaches for cleaning the Landsat data (Bell et al., 2023) and excluded those quarters of a year that had no data for more than 25% of the 30-m-grid pixels. For the first analysis we used data only from 2000 onwards because of limited image availability in some regions of Baja California due to fewer downlink stations for Landsat and storage issues during the 1980s and 1990s. For the precondition analysis in the three regions of interest, we selected the first year without any missing data in the time series, resulting in a continuous time series covering 1991–2023 for Ensenada region, 2000–2023 for Punta Eugenia region, and 1992–2023 for Bahía Asunción region.

### Exposure of kelp forests to MHWs

We estimated the exposure of kelp forests to MHWs, which are warm periods where temperature exceeds the 90^th^ percentile threshold relative to a baseline climatology (seasonal varying running mean) for at least five consecutive days (Hobday et al., 2016). We obtained daily SST estimates for 1982–2023 for our study region from the NOAA 0.25°-grid resolution OISST dataset (Huang et al., 2021), and identified pixels that overlaid with the remote sensed kelp-forest data. We then used the R package heatwaveR (Schlegel & Smit, 2018) to estimate the annual cumulative MHW intensities (°C days) registered from 2000–2023, relative to a 30-year baseline climatology (1983– 2012). Annual cumulative intensities are good indicators of species’ exposure to warm anomalies in a year (Arafeh-Dalmau et al., 2023; Oliver et al., 2019). We estimated mean annual cumulative MHW intensities across pixels for northern Baja California (n=16) and central Baja California (n=17).

### Underwater surveys of kelp, urchins, and urchin predators

To evaluate the potential mechanisms of kelp-forest loss and recovery after the MHWs we conducted field surveys of urchins, kelps, and urchin predators in 2022 and 2023. These surveys estimated *in situ* abundances of kelp, fish, and invertebrate species at 36 sites from Isla Coronado (32.5° N) near the USA-Mexico border to Bahía Asunción (27.1° N), representing 800 km of temperate transition zone along the coast of western Mexico (Figure 1). We followed the MasKelp protocol (similar to Reef Check (Freiwald et al., 2021) and PISCO (Malone et al., 2022) for underwater surveys) for Baja California, Mexico, which is a comprehensive protocol for surveying floating kelp forests based on remote-sensing and underwater surveys (Arafeh-Dalmau et al., 2019; García-Pantoja et al., 2025)

We used published maps of historical giant kelp-forest presence (Arafeh-Dalmau et al., 2021) for the region to select sites across the distribution of giant kelp forests in Mexico. We surveyed kelp forests that measured at least 1 km in length and selected two sub-sites within each forest. Sub-sites were selected near the center of the forest and were approximately 500 m apart from one another. In each sub-site, three 30 × 2 m transects were surveyed on scuba, parallel to the coastline and towards the direction of the other sub-site, for a total of six transects at each site. We quantified the abundance of kelp, macro-invertebrates (size > 2.5 cm), and fish species, as well as the size of all fish in 5-cm intervals.

### Compilation of existing long-term underwater surveys

We compiled existing long-term underwater datasets of urchins, giant and palm kelp to understand whether preconditions were predictive of the three alternative stable states observed in the region (i.e., giant-kelp forests, palm-kelp forests, or urchin barrens). For the Ensenada region that is today dominated by sea urchins (see Results section), we used six sites that have been surveyed from 2011–2013, 2016, and 2019–2021. Sites were surveyed by the Autonomous University of Baja California and The University of Queensland, following methods described in Arafeh-Dalmau et al. (2019), which are similar to those used in our 2022–2023 expeditions. The only difference is that those surveys used video rather than visual transects to estimate invertebrate abundances. We combined both datasets and developed a time series with seven sites from 2011–2023.

For the Punta Eugenia region that is today dominated by extensive giant kelp forests, we used 18 years of data from underwater surveys from Isla Natividad conducted annually by members of the fishing community with the support of Comunidad y Biodiversidad A.C. and Stanford University, following Reef Check protocol (Freiwald et al., 2021) between July–September from 2006–2023 within five sites. The protocol is very similar to MasKelp, the main difference being the number of transects and the way sites are selected. At each site, 14–27 transects were surveyed yearly (av. = 19.93, SE = 0.40) in shallow (average 7-m depth) and deeper waters (average 14 m).

For the southern region at the distribution limit of giant kelp that is today dominated by extensive palm-kelp forests, we used 21 years of underwater surveys at six sites (Chávez-Hidalgo, 2017). The six sites from San Pablo to Isla San Roque (∼27.1° N) were surveyed annually (autumn) from 1993–2013 by members of the fishing cooperative and the Instituto Nacional de Pesca (INAPESCA) to a maximum depth of 23 m. We also included density estimates for palm kelp from 2019 and the 2022–2023 expeditions conducted at two sites (six transects/site). We included only urchin density estimates that were available for 2022–2023. See Table S1 for all sites and years included.

All sites that fell in the same 0.5°-latitudinal bin were used to aggregate the remote-sensed giant-kelp dataset for the three regions and allowed us to create a long-term time series of urchin and palm-kelp densities, and giant-kelp coverage. We chose to include giant-kelp area estimates from remote sensing because this data is the most comprehensive long-term timeseries available in Mexico, and because density estimates from underwater surveys are not available in Bahía Asunción before the MHWs.

### Sea surface temperature and human gravity index

To evaluate whether SST and human impact may be drivers of the three alternative states in northern and central Baja California, we obtained daily SST data from the NOAA-ERDDAP data repository (https://coastwatch.pfeg.noaa.gov/erddap/index.html), for the years post-MHWs (2017–2023) from the Aqua-MODIS satellite at 1-km^2^ grid and estimated the mean annual SST post-MHWs for each of the 72 sub-sites surveyed in 2022 and 2023. Then, to estimate a metric of human impact along the coastline of our region, we estimated the human gravity index, following the approach of Cinner et al. (2018).This index captures the intensity of human presence by integrating both the population size and proximity of nearby coastal settlements, providing a proxy for the potential exposure of each kelp pixel to human impact. We used publicly available datasets on settlement location and population size from the 2020 national census in Mexico (INEGI; https://www.inegi.org.mx/programas/ccpv/2020/#datos_abiertos) and the 2020 American Community Survey in the United States (https://www.inegi.org.mx/app/descarga/ficha.html?tit=326108&ag=0&f=csv). We retained only settlements within 50 km of the coastline. We calculated the human gravity index on the same 1-km^2^ SST pixels using a distance-weighted function that incorporates both settlement size and distance.

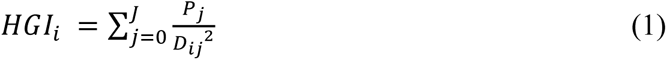

where HGI is the human gravity index of kelp pixel *i* summed across all human settlements *j* located within a 50 km radius of kelp pixel *i*. For each human settlement *j*, *P_j_* is the population size, and *D_ij_* is the distance between kelp pixel *i* and settlement *j*.

### Data analysis

We used generalized additive mixed models (GAMMs) with a Tweedie distribution and a log link function to model remote-sensed kelp canopy area as a function of year (as a smooth term) and ecoregion (northern or central Baja California), including their interaction. We included a random intercept for each 1-km^2^ pixel to account for repeated measures in time. We used generalized mixed effects models (GLMMs) with a negative binomial distribution and a log link function to accommodate overdispersion to model count data on the abundance of giant and palm kelp and lobsters as a function of ecoregion and year, including their interaction. For urchin abundance and sheepshead biomass we used a Tweedie distribution and a log link function because corresponding AIC and residuals were better. For the GAMM and the GLMMs, we also included a spatial random field by constructing a mesh based on the geographic coordinates of the survey sites. When fitting these models, we estimated the Tweedie power parameter jointly with the model coefficients. For all GLMM we included a random intercept for each site to account for repeated measures. More complex random effect structures (e.g., transects nested within sites) led to convergence issues and were therefore excluded.

We then simplified models by removing the interactions and terms if they were not significant (p ≤ 0.05) and their removal did not cause a deterioration in model fit (AIC). For giant kelp and urchins, we retained the full model, for sheepshead we removed the ecosystem-by-year interaction, and for palm kelp and lobster we removed both the interaction and the term for year. Additionally, we used GLMMs to model results of long-term underwater surveys for palm kelp and urchins for Ensenada and Punta Eugenia region as a function of heatwave period (before and after) and included random intercepts for sites to account for repeated measures. For Punta Eugenia region, we included temporal autocorrelation using an autoregressive correlation structure (AR1) grouped by site. For Ensenada region we included year as a random intercept because we had many gaps in the time series. For this analysis, the limited number of sites meant that we could not reliably adjust for spatial autocorrelation.

We fit GLMMs with spatial mesh using the “sdmTMB” package (Anderson et al., 2022), assessed statistical significance of fixed effects using 95% confidence intervals (CIs) derived from model standard errors, and considered effects significant when CIs did not include zero. We fit models without spatial mesh using the “glmTMB”package (Brooks et al., 2017) and “car” to compute Wald Tests of the main effects (Type III). We evaluated residual diagnostics for all models using “DHARMa” (Hartig, 2018); associated plots are presented in the Supplemental Material (Figure S1–S10).

We used nonlinear quantile regression of the median abundances (using the R package *quantreg (Koenker, 2023)*) to assess the relationships of the trophic cascade between kelp, urchins, and their predators in Baja California, and for the relationship between urchin abundance and sheephead biomass. In addition, we assessed the relationships between giant kelp and palm kelp species to assess whether competition among species is a driver of kelp change in Baja California. We used quantile regressions to characterize the boundary conditions of relationships, and we standardized on the 88^th^ percentile because higher percentiles resulted in model instability for at least one of the pairs of predictors.

To model the probability of species dominance as a function of climate and human impact, we fit a multinomial logistic regression with regime state (giant kelp, palm kelp, or urchin-dominated) as a categorical response variable. We derived biomass values from mean biomass estimates obtained from Woodson et al. (2019) and assigned dominance state at each site to the taxon with the greatest estimated biomass. Predictors included mean annual SST after the MHWs (2017– 2023) and the human gravity index. We used giant kelp as the reference category. Model outputs were used to generate predicted probabilities across the SST and human gravity index. We assigned statistical significance using Wald tests.

To detect outliers in the analysis of the Landsat-based time series of kelp area, we used automatic algorithms from the R package *tsoutliers (López-de-Lacalle, 2016)* to fit ARIMA (0,1,1) time series to the data for each region. This method offers an advantage over random-walk models by smoothing the fit using an exponentially weighted moving average of prior values, rather than placing excessive emphasis on the most recent observation. Outliers identified in this way can be considered as arising from processes external to the natural variation in the time series. The main types of outliers that can be detected include additive outliers, which represent sudden, one-time changes affecting only a single observation without impacting the rest of the time series; temporary changes, which involve smooth transitions that eventually revert to earlier conditions; and level shifts, which indicate lasting changes in the mean that persist over time.

All analyses and data preparation were conducted in R.

## Results

### Kelp coverage and marine heatwave intensities

Giant kelp forests declined across Baja California following the MHWs, except in the region of Punta Eugenia in central Baja California, where kelp coverage increased (Figure 1, Figure S11). We documented a range contraction of half a degree of latitude at the southern (equatorward) distribution limit of giant kelp (∼99.9% loss in the area spanning 27–27.5° N), equivalent to a straight-line distance of ∼80 km, where giant kelp has been absent since 2015 (Figure 1, Figure S11). We also observed loss of kelp along approximately 0.9° of latitude (∼110 km) in the northern part of our study area, from the USA–Mexico border to Punta Banda, where giant kelp has been absent since 2020, corresponding to an estimated 99% loss. These observed shifts in range are supported by underwater surveys conducted in 2022 and 2023, during which we did not find giant kelp at the three southernmost and eight northernmost monitoring sites (see next sections).

From 2000–2013, giant kelp forests in Mexico covered an average of 11.46 km² annually, spanning 1,605.7 km of island and continental coastline across 5.5° of latitude (Figure 1). During the 2014– 2016 MHWs, kelp forests were exposed to an average cumulative annual intensity of 528.4 ± 47.9 °C days—an order of magnitude higher than the 29.3 ± 4.2 °C days recorded during the pre-MHWs period (Figure 2a). As a result, kelp cover across Baja California declined by 45.1% during the MHWs (Figure 1, Figure 2). The most extreme year was 2015, with intensities reaching 821.0 ± 30.5 °C days, and by 2016, only 1.33 km² of kelp remained—an 88.4% reduction from the pre-MHWs baseline.

**Figure 2.**
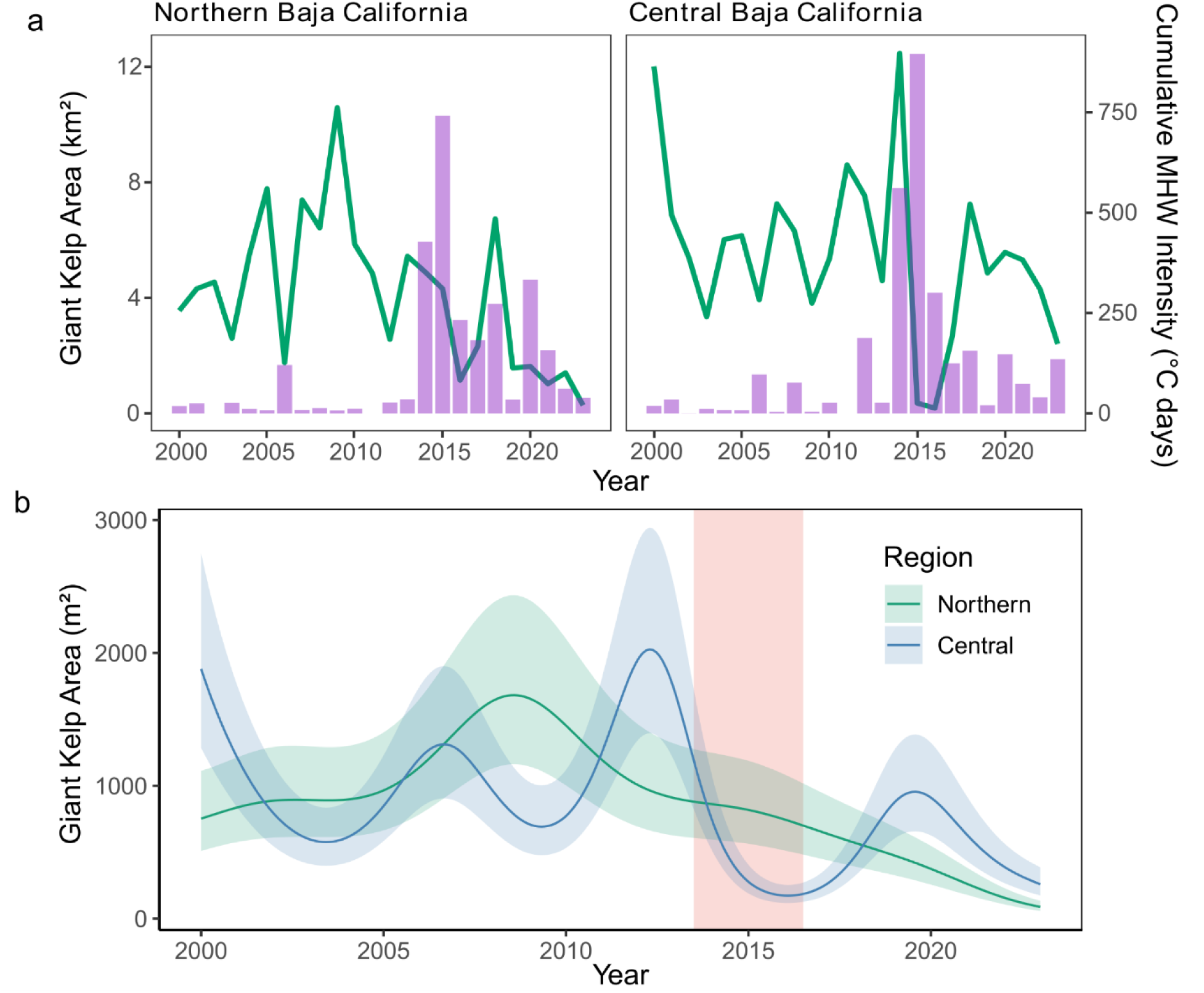
- Response of giant kelp forests to the 2014-2016 MHWs. Panel **a** is the yearly mean kelp area (green line, left y-axis) plotted with the mean annual cumulative MHW intensities per 0.25° pixel for northern (n=16) and central (n=17) Baja California ecoregions (purple bar plots, right y-axis). Pannel **b** illustrates estimated mean kelp area per 1 km² pixel for northern (n=470, cyan color) and for central (n = 416, blue color) Baja California ecoregions from temporal generalized additive mixed models (GAMMs) using Gaussian-process splines. Lines represent predicted values, and shaded bands show 95% confidence intervals. Pink shading indicates the 2014–2016 MHW period.

MHW exposure and kelp response varied regionally. During the MHWs, central Baja California experienced higher cumulative annual intensities (585.8 ± 70.1 °C days) than northern Baja California (467.4 ± 60.9 °C days), with corresponding kelp losses of 61.9% and 20.8%, respectively (Figure 2, Table S2). After the MHWs (2017–2023), cumulative annual intensities remained elevated (126.9 ± 92.7 °C days) but were higher in northern Baja California (154.2 ± 22.0 °C days) than central Baja California (99.6 ± 11.7°C days). The GAMM output (Figure 2b) captures these trends, showing a pronounced initial decline in kelp cover in central Baja California, followed by partial recovery after 2017. In contrast, northern Baja California exhibited a less pronounced initial decline but with a continuous downward trajectory post-MHWs.

Since 2017, kelp cover has fluctuated, with a peak above mean pre-MHWs levels in 2018 in both regions—suggesting initial recovery. However, this trend was short-lived. Northern Baja experienced steady declines from 2021 onward, reaching over 90% loss by 2023 (Figure 2a). Central Baja showed more variable patterns, with the steepest decline in 2023 (61.8%; Figure 2b), likely driven by Hurricane Hilary, a Category-4 hurricane that made landfall in August 2023.

### Mechanisms of giant kelp recovery and loss

Underwater surveys corroborated the remote-sensing evidence of giant kelp range contraction, including the loss of over 100 km of kelp coverage near the USA–Mexico border and complete absence of kelp in the southern range. Giant kelp was absent at the three southernmost sites (n = 18 transects per year) and at the eight northernmost sites across both years (n = 48 transects per year). In contrast, kelp abundances peaked in central Baja California, where densities of giant and palm kelp were 36 and 62.1-fold higher, respectively, than in northern Baja California (*p* < 0.005, Table S3-S4). Urchin predators showed similar spatial trends, with lobster abundance (*Panulirus interruptus*) and California sheephead (*Semicossyphus pulcher*) biomass 18.3 and 16.3 times higher, respectively, in the central region compared to the north (*p* < 0.0001). Predators were nearly absent north of 29°N, where only five lobsters were recorded across 148 transects surveyed during 2022 and 2023 (Figure 3a). In contrast, sea urchin densities were significantly higher in northern Baja California, with abundance 23.2 times greater than in the central region (*p* < 0.0001). Three herbivore urchin species—*Strongylocentrotus purpuratus*, *Mesocentrotus franciscanus*, and *Centrostephanus coronatus*—were present in both regions.

**Figure 3.**
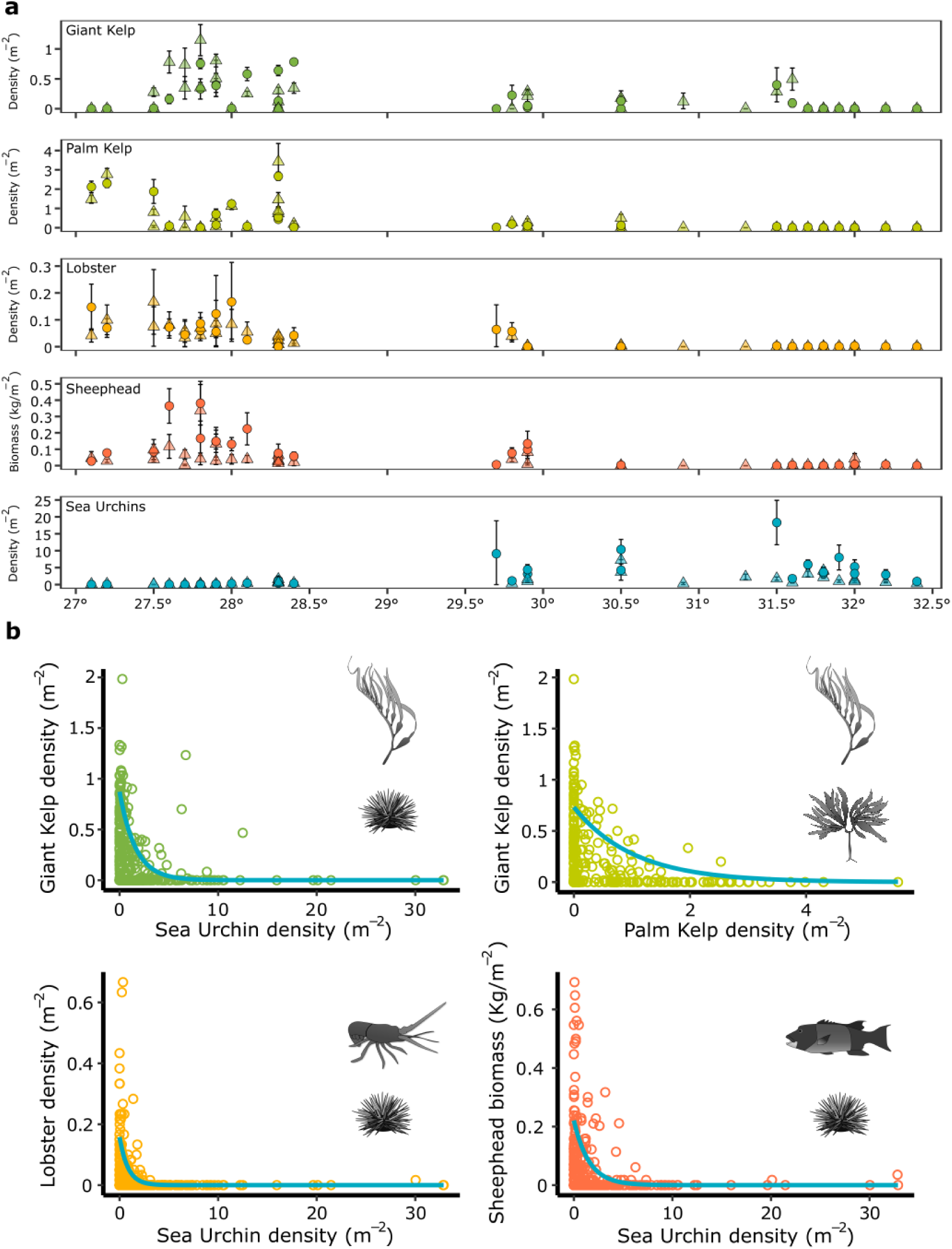
- Mechanisms driving giant kelp recovery and loss from MHWs. Panel **a** is the latitudinal density of kelps, urchin predators, and sea urchins (mean number of individuals m^-2^, n = two sub-sites with three census transects in each, ±95% confidence intervals) at 35 and 32 sites in 2022 (triangles) and 2023 (circles), respectively. For sheephead, we estimated the biomass of large individuals (size > 35 cm total length) that can consume both juvenile and adult urchins. Panel **b** is the relationship among kelps, urchins, and urchin predators from a non-linear quantile regression of the 88^th^ percentile, using the data from panel **a**.

When we used quantile regression to assess the relationships between kelp, urchins, and their predators in Baja California, we found a negative relationship between the densities of giant kelp and urchins (Figure 3b; *p* < 0.00001). Urchin densities were also lower with higher densities of both California sheephead (*p*: 0.00024) and lobster (*p* < 0.00001), the main predators of urchins in these ecosystems. Finally, we found a significant negative relationship between the abundance of giant and palm kelp (p: 0.00035), suggesting competition between these species. Taken together, these results reveal three alternative states in Baja California: sea urchin barrens in most of northern Baja California; giant kelp forests, mainly in parts of central Baja California; and palm kelp forests in other parts of central Baja California.

Multinomial logistic regression confirmed strong and opposing effects of SST on state probability: SST was positively associated with palm kelp (β = 5.35, *p* < 0.0001) and negatively with urchins (β = –3.88, *p* < 0.0001), both relative to giant kelp. Human impact significantly increased the likelihood of urchin dominance (β = 0.002, *p* = 0.044) but had no detectable effect on palm kelp (β = -0.016, *p* = 0.21). Modeled species dominance aligned with gradients in sea surface temperature (SST) and human impact (Figure 4): urchin-dominated states were most probable under cooler SST and high impact; giant kelp under intermediate SST and low to moderate impact, and palm kelp under the warmest conditions. These patterns highlight the combined climate-and impact-driven restructuring of coastal ecosystems, with kelp- and predator-dominated communities more likely in the warmer, less impacted south, and urchin-dominated states prevailing in the cooler, more impacted north. Notably, giant kelp remained dominant even at sites where maximum monthly SSTs after the MHWs exceeded 22.5°C (Figure S12).

**Figure 4.**
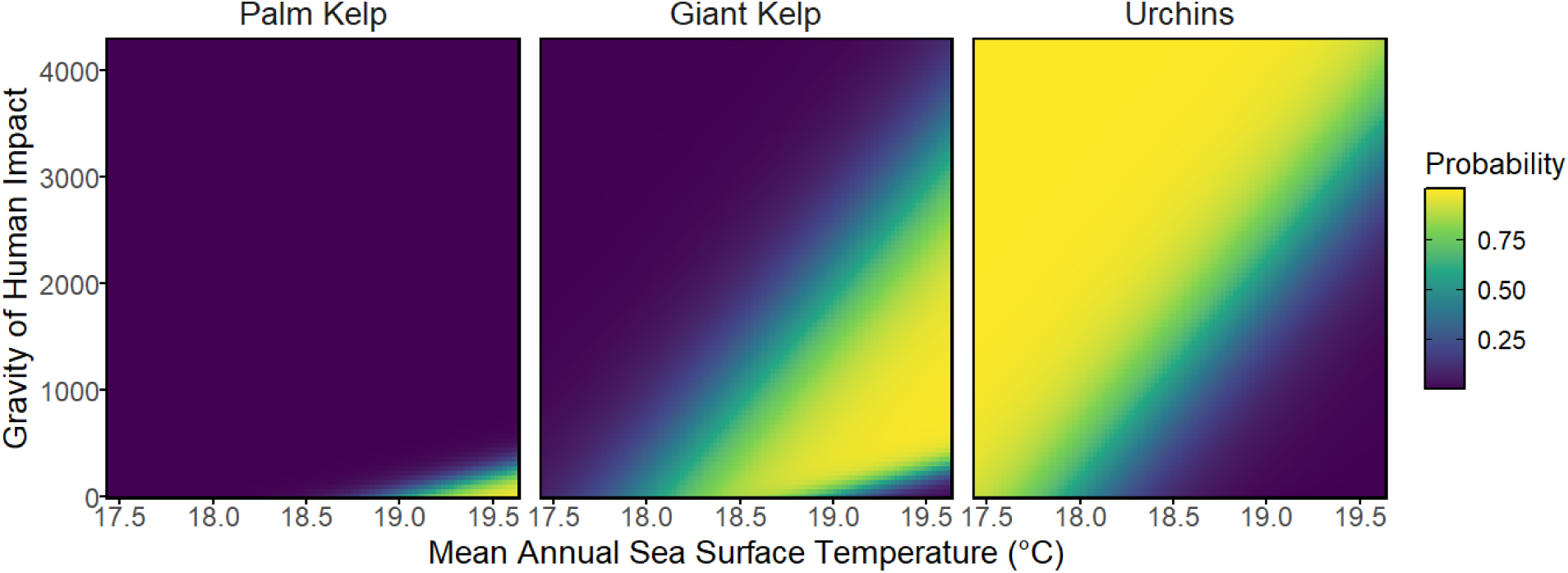
- Predicted probabilities of alternative ecological states across gradients of temperature and human impact. Each panel shows the modeled probability of dominance for palm kelp, giant kelp, and urchins across 72 sub-site observations (36 sites with two sub-sites) aggregated for 2022– 2023 in the Baja California Peninsula, Mexico. Probabilities are derived from a multinomial model incorporating environmental predictors, with outputs visualized across the environmental space. Mean annual sea surface temperature (°C) (after MHWs: 2017-2023) and gravity of human impact are continuous variables, while color gradients indicate the relative probability of each state, ranging from low (purple) to high (yellow).

### Preconditions driving alternative stable states

Using historical satellite and underwater survey data, we assessed kelp trajectories and ecological preconditions prior to recent MHWs across three Mexican regions. ARIMA analysis identified 16 significant outliers in kelp canopy area in the Ensenada region (northern Baja California; 31.75-32.25°) between 1991 and 2023, indicating long-term instability. Most outliers were additive, reflecting abrupt, one-time changes or temporary changes that gradually returned to prior levels (Figure 5, Table S5). From 1991–2007, kelp cover remained relatively low and stable (four outliers), followed by increased variability and higher coverage from 2008 to 2013 (six outliers). During and after the 2014–2016 MHWs, two level shifts were detected, reflecting persistent changes in the mean: a major decline in 2015, followed by a brief rebound in 2017. This was succeeded by a sustained decline through 2023, resulting in a 90% kelp loss by the end of the time series (Figure S11). In contrast, Punta Eugenia (central Baja California; 27.5-28°) showed stable kelp canopy from 2000 to 2023, with only two additive outliers and no level shifts detected during the MHWs (Table S6). Bahía Asunción (central Baja California; 27-27.5°) exhibited an intermediate pattern, with five outliers including four level shifts (Table S7). Three distinct canopy states were evident: low coverage (1993–1999), high and stable coverage (2000–2014), and persistent collapse beginning in 2015, with >99% kelp loss.

**Figure 5.**
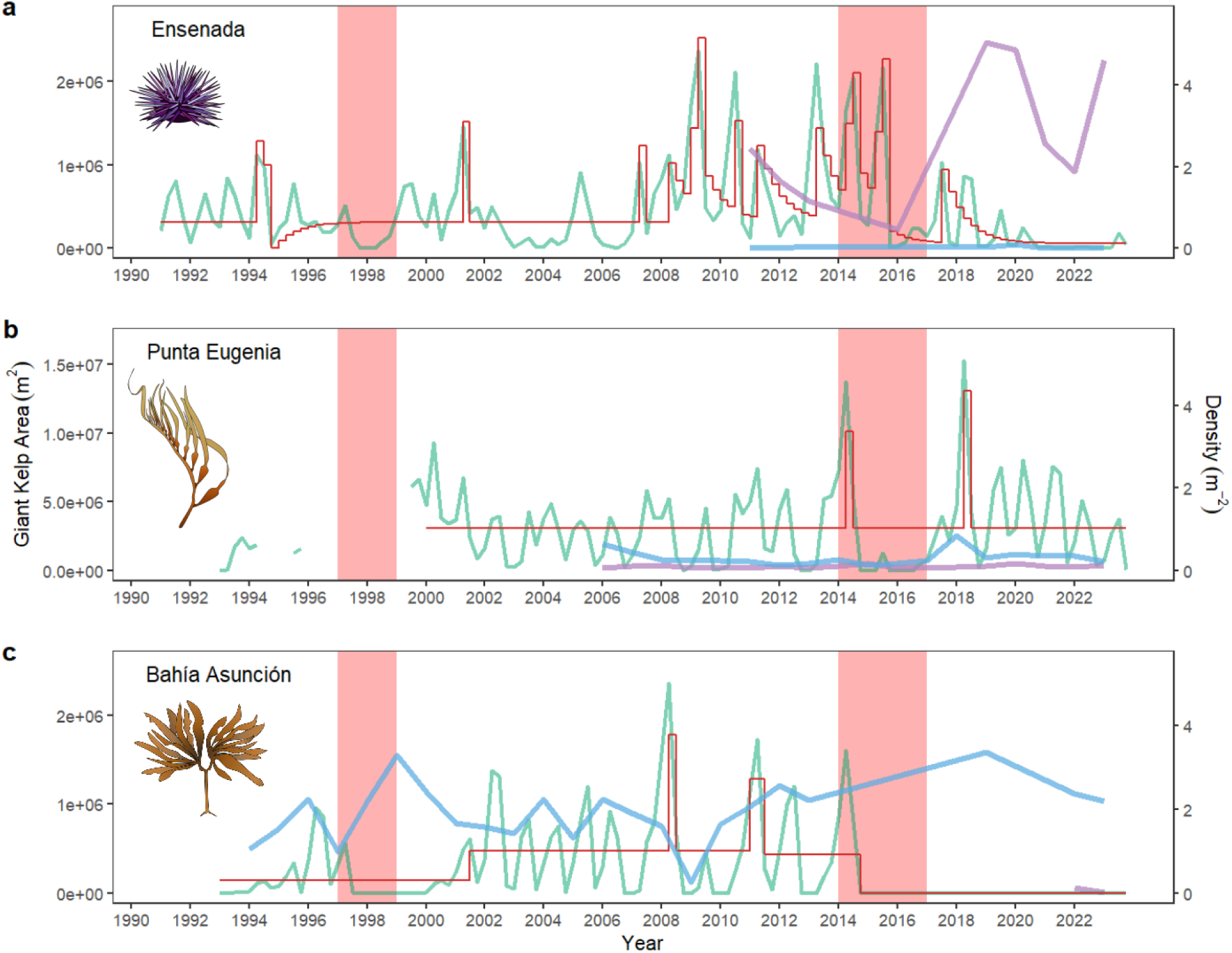
- Preconditions driving alternative states and the recovery or loss of giant kelp from MHWs. Time series of remote-sensed kelp area (m^2^, green lines) aggregated at 0.5° of latitude for the Ensenada (**a**, 1991–2023, 31.75–32.25° N), Punta Eugenia (**b**, 2000–2022; 27.5–28° N), and Bahía Asunción (**c**, 1993–2023; 27–27.5° N) regions. Superimposed on each kelp time series (in red) is the estimated deviation from the expectation of an ARIMA (0,1,1) fitted area model. Purple and blue lines represent time series of urchin and palm kelp densities, respectively (mean number of individuals m^-2^, ±95% confidence intervals), for each region. Note that for Bahía Asunción we only have the mean estimates for palm kelp. The vertical red bars correspond to the 1997-98 ENSO event and the 2014-16 MHWs. Note that the y-axis scales for Ensenada and Bahía Asunción differ from Punta Eugenia.

Ecological preconditions differed across regions and likely influenced these divergent kelp trajectories after the MHWs (Figure 5). In Ensenada, sea urchin densities were relatively high before the MHWs (2011–2013: 1.26 ± 0.44 individuals m^-2^) and increased significantly afterward (2019–2023: 3.05 ± 0.92 individuals m^-2^; p = 0.0029), consistent with grazer-driven collapse of kelp forests. Palm kelps were rare and did not significantly increase after the MHWs (<0.005 individuals m^-2^; p = 0.533). In Punta Eugenia, both urchins (<0.08 individuals m^-2^: p = 0.5724) and palm kelps (<0.35 individuals m^-2^; p = 0.3354) remained at low, stable densities before and after the MHWs, consistent with a stable giant kelp ecosystem. In Bahía Asunción, at the historical southernmost limit for giant kelp, palm kelp densities were substantially higher, increasing from 1.83 ± 0.67 individuals m^-2^ before the MHWs (1994–2013) to 2.56 ± 0.69 individuals m^-2^ afterward (2019, 2022–2023). During the 1997–1998 El Niño, palm kelp peaked at 3.3 individuals m^-2^, while giant kelp was absent until 2000–2001, after which palm kelp declined, and giant kelp recovered — suggesting competitive interactions likely limited giant kelp reestablishment following the 2014–2016 MHWs. Urchin data were unavailable for Bahía Asunción prior to the MHWs, but post-MHW densities were low (0.06 ± 0.07 individuals m^-2^ in 2022–2023).

Pre-MHW urchin densities were 19.4 times higher in Ensenada than in Punta Eugenia, and palm kelp densities in Bahía Asunción were 5.7 and 68.3 times higher than in Punta Eugenia and Ensenada, respectively. We report raw means for comparability across regions because model estimates were not available for Bahía Asunción. Together, these patterns suggest that divergent post-MHW kelp forest trajectories were mediated by ecological preconditions — particularly grazer pressure and the competitive dominance of palm kelp — which mediated whether giant kelp remained stable or transitioned to an alternative state.

## Discussion

Four decades of remote sensing and field observations reveal that extreme MHWs can restructure ecosystems by driving regime shifts and multiple alternative states. We document an 80 km southern (equatorward) range-edge contraction and the emergence of three distinct ecosystem states near the warm limit of giant kelp forests in Mexico. Before the 2014–2016 MHWs, all these systems were dominated by giant kelp, but post-MHW, trajectories diverged based on local conditions. In central Baja California, warm-tolerant palm kelp (Almeida-Saá et al., 2025; Edwards & Hernandez-Carmona, 2005) displaced giant kelp in some areas, while most giant kelp remained resilient. Unexpectedly, the most severe losses occurred in colder northern regions, where predator depletion enabled urchin overgrazing and widespread kelp collapse. Along a 110-km stretch, 99% of kelp forests have been lost and substituted by urchin barrens, likely triggering cascading impacts on biodiversity, ecosystem services, and fisheries (Arafeh-Dalmau et al., 2019; Smith et al., 2021; Villaseñor-Derbez et al., 2024). These findings challenge expectations of warming-driven decline near equatorward limits (Beas-Luna et al., 2020) and show that while MHWs can drive ecosystem collapse (McPherson et al., 2021; Smith et al., 2023; Wernberg et al., 2016), the direction and extent of change depend on pre-existing conditions (J. G. Smith et al., 2024) —such as high predator biomass—that can buffer climate impacts (Kumagai et al., 2024). By documenting three coexisting stable states and their climatic and ecological drivers, we contribute new insights into climate-driven regime shifts and the role of species interactions in ecological resilience.

The documentation of three coexisting stable states within a climate-change hotspot (Hobday & Pecl, 2014) is unprecedented. While regime shifts in kelp forests are usually binary—transitioning either to urchin barrens (McPherson et al., 2021; Rogers-Bennett & Catton, 2019; J. G. Smith et al., 2024) or turf-dominated systems (Filbee-Dexter et al., 2020; Wernberg, 2021; Wernberg et al., 2016)—we demonstrate the co-existence of a third distinct state: domination by the smaller palm kelp. This state has persisted at the southernmost sites for over nine years in Baja California Sur since the 2014–2016 MHWs, in contrast to the 1999 El Niño, when giant kelp forests were temporarily displaced and replaced by palm kelp, but fully recovered within two years (Edwards & Hernandez-Carmona, 2005). As MHWs intensify (Arafeh-Dalmau et al., 2025), giant kelp and other cold-affinity foundation species may increasingly be displaced by more thermally tolerant taxa, particularly near the warm limits of their distributions (Arafeh-Dalmau et al., 2019; Filbee-Dexter et al., 2020; Wernberg, 2021). While prior models align with this expectation (Beas-Luna et al., 2020), our empirical observations challenge these forecasts: resilient and dense giant kelp forests can also persist near their southern range limit. This resilience is likely due to low human pressure (Donovan et al., 2021), effective management (Micheli et al., 2012; Olguín-Jacobson et al., 2025), the presence of local climate refugia (Arafeh-Dalmau et al., 2021) (i.e., areas where local conditions buffer climate change impacts), population connectivity (Vranken et al., 2025), and potential local adaptation (Torda et al., 2017; Wernberg et al., 2018)—factors often overlooked in predictive models.

A central finding is that preserving intact populations of top predators prevents undesirable regime shifts in marine ecosystems, suggesting their conservation can be an effective climate-adaptation strategy. The collapse of kelp forests and emergence of extensive urchin barrens in northern Baja California coincided with the near-complete absence of predators, while central Baja California ecosystems with high predator abundance returned to a state of kelp dominance relatively quickly. Overfishing is likely undermining trophic integrity and increasing the risk of climate-driven collapse (Kumagai et al., 2024; Ling et al., 2009; Ling & Keane, 2024). Similar patterns were observed during the 2014–2016 MHWs in southern California, where trophic cascades supported the resilience of kelp forests in a network of 39 marine protected areas (Kumagai et al., 2024). Our findings suggest that this mechanism also operates in central Baja California, where kelp forests are effectively managed by fishing cooperatives through community-based marine reserves, catch and size limits, and seasonal closures (McCay et al., 2014; Micheli et al., 2024; Micheli et al., 2012; Olguín-Jacobson et al., 2025). These results suggest that locally driven governance schemes and stewardship such as “other effective area-based conservation measures” (OEBCMs) can be an additional climate adaptation tool to enhance ecosystem resilience in the face of intensifying climate impacts and advance global 30 x 30 biodiversity targets (Bennett et al., 2022; CBD, 2022).

Long-term remote sensing data and underwater surveys reveal that regime-shift trajectories are strongly shaped by pre-disturbance dynamics (J. G. Smith et al., 2024). Giant kelp forests with low grazer and competitor densities were stable and recovered post-heatwaves, while those with prior variability or collapse were more likely to shift into urchin barrens or palm kelp dominance. Sea surface temperature and human impact explain these alternative states: giant kelp dominates across a range of temperatures when human pressure is low; urchin barrens emerge in cooler, more impacted areas; and palm kelp dominates warmer, less disturbed sites. Shifts to urchins were gradual, while palm kelp transitions were abrupt, indicating nonlinear ecological responses. Notably, high temporal variability before disturbance may signal vulnerability (Oliver et al., 2015), highlighting stable areas that are more likely to persist as conservation priorities, as well as early interventions in variable vulnerable areas. These findings highlight the importance of a portfolio of climate adaptation strategies, including monitoring ecosystem stability (White et al., 2025), predator protection, grazer control(Eger et al., 2022), managing novel ecosystems (Bay et al., 2023; Webster et al., 2023), or mitigating human impacts (Arafeh-Dalmau et al., 2023; Kumagai et al., 2024) to enhance resilience under intensifying climate extremes (Arafeh-Dalmau et al., 2025; IPCC, 2023; Oliver et al., 2019).

The scarcity of long-term *in situ* data limits our understanding of the mechanisms driving observed regime shifts. Subtidal surveys in northern Baja California were spatially broad but temporally inconsistent, while in central Baja California, longer-term data were restricted to a few locations and inconsistently sampled in some regions. Additionally, differences in sampling methods also limited regional comparisons and hindered understanding the role of urchin predators during and after the heatwaves. Our 2022–2023 expeditions addressed some of these gaps by simultaneously and consistently surveying 36 sites across the full latitudinal range of giant kelp in Mexico, enabling integration of historical and contemporary data. This synthesis confirms that trophic cascades and species competition are key drivers of post-heatwave ecosystem dynamics, consistent with ecological theory. These limitations highlight the urgent need for standardized, long-term monitoring of vulnerable coastal ecosystems to detect early warning signals, improve mechanistic understanding, and guide adaptive responses to climate-driven ecosystem change (Hughes et al., 2017; Kumagai et al., 2024; Olguín-Jacobson et al., 2025; Schmeller et al., 2018).

Our findings show that MHWs can drive regime shifts, but the trajectories of change are shaped by ecological preconditions. We observed three persistent ecosystem states along gradients of temperature, grazing pressure, and human impact. Remarkably, giant kelp persisted in central Baja California despite mean annual maximum monthly temperatures exceeding ∼22.5 °C —conditions that are likely near or beyond physiological thresholds (Cavanaugh et al., 2019; Harden et al., 2024; Le et al., 2022; Rothäusler et al., 2011) for giant kelp. This resilience likely reflects a combination of ecological integrity, local adaptation, and effective management. In contrast, forests in cooler, more impacted northern regions collapsed where predators were absent, and herbivores proliferated. These outcomes highlight the need to integrate biotic interactions and management context into climate adaptation strategies. Protecting ecologically intact, heat-tolerant ecosystems, especially through community-based management, may be key to enhancing resilience, preventing regime shifts, and support biodiversity and coastal livelihoods under a warming ocean.

## Acknowledgments

N.A.-D., F.M., K.C., and G.M-M, acknowledge funding from the Lenfest Ocean Program (ID Number: 00036969). N.A.-D. from the Estate Winifred Violet Scott (Australia) to conduct field expeditions. F.M., C.O-J., and N.A-D. from the National Science Foundation (2108566). K.C. from NASA Ocean Biology and Biogeochemistry program (80NSSC21K1429) and the National Science Foundation SBC LTER Award (1232779). G.M-M. from UABC (ID Number: 3916) and SEMARNAT-INE-CONACYT 2008/01-Fondo Sectorial de Investigación Ambiental. A.M.-V. from the National Geographic Society via the Exploration Technology Lab Deep-Sea Collaboration. O.A.-O., from the Baum Foundation, to support expeditions. We are very thankful to the Federation of Fishing Cooperatives of Baja California and Coperativa Ensenada, Sirenas del Mar, in addition to all the fishers, divers, and institutions that have supported field expeditions.

## Author Contributions Statement

N.A-D. conceived the study with inputs from F.M., K.C., and D.S. N.A-D. led field expeditions with the support of A.M-V., C.O-J., O.A-O., F.M., G.M-M., and J.G-P. N.A-D., C.O-J., J.G-P., G.T-M. collected data. N.A.-D conducted analyses with the support of D.S. N.A-D. led the writing of the manuscript with the support of D.S, F.M., and K.C. A.M-V., C.O-J., O.A-O., J.G-P., G.M-M, and G.T-M. contributed to reviewing and editing of the manuscript.

## Data Availability

The remote-sensing kelp forest dataset is available at https://portal.edirepository.org/nis/mapbrowse?packageid=knb-lter-sbc.74.13. All other data needed to evaluate the conclusions in the paper are present in the paper or its Supplementary information. The kelp forest underwater surveys will be made available upon publication.

## Code Availability

The codes used for this project will be made available upon publication.

## References

Almeida-Saá, A. C., Umanzor, S., Zertuche-González, J. A., Cruz-López, R., Ferreira-Arrieta, A., Rangel-Mendoza, L. K., & Sandoval-Gil, J. M. (2025). Comparative photophysiology and respiration of Kelp gametophytes reveals species-specific thermo-tolerance to marine heatwaves. Marine Biology, 172(5), 71.

Anderson, S. C., Ward, E. J., English, P. A., & Barnett, L. A. (2022). sdmTMB: an R package for fast, flexible, and user-friendly generalized linear mixed effects models with spatial and spatiotemporal random fields. bioRxiv, 2022.2003. 2024.485545.

Arafeh-Dalmau, N., Cavanaugh, K. C., Possingham, H. P., Munguia-Vega, A., Montaño-Moctezuma, G., Bell, T. W., Cavanaugh, K., & Micheli, F. (2021). Southward decrease in the protection of persistent giant kelp forests in the northeast Pacific. Communications Earth & Environment, 2(1), 1–7.

Arafeh-Dalmau, N., Montaño-Moctezuma, G., Martínez, J. A., Beas-Luna, R., Schoeman, D. S., & Torres-Moye, G. (2019). Extreme marine heatwaves alter kelp forest community near its equatorward distribution limit. Frontiers in Marine Science, 6, 499.

Arafeh-Dalmau, N., Munguia-Vega, A., Micheli, F., Vilalta-Navas, A., Villaseñor-Derbez, J. C., Précoma-de la Mora, M., Schoeman, D. S., Medellín-Ortíz, A., Cavanaugh, K. C., & Sosa-Nishizaki, O. (2023). Integrating climate adaptation and transboundary management: Guidelines for designing climate-smart marine protected areas. One Earth, 6(11), 1523–1541.

Arafeh-Dalmau, N., Schoeman, D. S., Montaño-Moctezuma, G., Micheli, F., Rogers-Bennett, L., Olguin-Jacobson, C., & Possingham, H. P. (2020). Marine heat waves threaten kelp forests. Science, 367(6478), 635–635.

Arafeh-Dalmau, N., Villaseñor-Derbez, J. C., Schoeman, D. S., Mora-Soto, A., Bell, T. W., Butler, C. L., Costa, M., Dunga, L. V., Houskeeper, H. F., & Lagger, C. (2025). Global floating kelp forests have limited protection despite intensifying marine heatwave threats. Nature Communications, 16(1), 3173.

Armstrong McKay, D. I., Staal, A., Abrams, J. F., Winkelmann, R., Sakschewski, B., Loriani, S., Fetzer, I., Cornell, S. E., Rockström, J., & Lenton, T. M. (2022). Exceeding 1.5 C global warming could trigger multiple climate tipping points. Science, 377(6611), eabn7950.

Baum, J. K., Claar, D. C., Tietjen, K. L., Magel, J. M., Maucieri, D. G., Cobb, K. M., & McDevitt-Irwin, J. M. (2023). Transformation of coral communities subjected to an unprecedented heatwave is modulated by local disturbance. Science Advances, 9(14), eabq5615.

Bay, L. K., Gilmour, J., Muir, B., & Hardisty, P. E. (2023). Management approaches to conserve Australia’s marine ecosystem under climate change. Science, 381(6658), 631–636.

Beas-Luna, R., Micheli, F., Woodson, C. B., Carr, M., Malone, D., Torre, J., Boch, C., Caselle, J. E., Edwards, M., Freiwald, J., Hamilton, S. L., Hernandez, A., Konar, B., Kroeker, K. J., Lorda, J., Montaño-Moctezuma, G., & Torres-Moye, G. (2020). Geographic variation in responses of kelp forest communities of the California Current to recent climatic changes. Global Change Biology, 26(11), 6457–6473. 10.1111/gcb.15273

Bell, T. W., Cavanaugh, K. C., Saccomanno, V. R., Cavanaugh, K. C., Houskeeper, H. F., Eddy, N., Schuetzenmeister, F., Rindlaub, N., & Gleason, M. (2023). Kelpwatch: A new visualization and analysis tool to explore kelp canopy dynamics reveals variable response to and recovery from marine heatwaves. PLoS One, 18(3), e0271477. 10.1371/journal.pone.0271477

Bennett, N. J., Le Billon, P., Belhabib, D., & Satizábal, P. (2022). Local marine stewardship and ocean defenders. NPJ Ocean Sustainability, 1(1), 3.

Brooks, M. E., Kristensen, K., Van Benthem, K. J., Magnusson, A., Berg, C. W., Nielsen, A., Skaug, H. J., Machler, M., & Bolker, B. M. (2017). glmmTMB balances speed and flexibility among packages for zero-inflated generalized linear mixed modeling. The R journal, 9(2), 378–400.

Cavanaugh, K. C., Bell, T., Costa, M., Eddy, N. E., Gendall, L., Gleason, M. G., Hessing-Lewis, M., Martone, R., McPherson, M., & Pontier, O. (2021). A Review of the Opportunities and Challenges for Using Remote Sensing for Management of Surface-Canopy Forming Kelps. Frontiers in Marine Science, 1536.

Cavanaugh, K. C., Reed, D. C., Bell, T. W., Castorani, M. C., & Beas-Luna, R. (2019). Spatial variability in the resistance and resilience of giant kelp in southern and Baja California to a multiyear heatwave. Frontiers in Marine Science, 6, 413.

CBD. (2022). COP15: Kunming-Montreal Global biodiversity framework. https://chrome-extension://efaidnbmnnnibpcajpcglclefindmkaj/https://www.cbd.int/doc/c/e6d3/cd1d/daf663719a03902a9b116c34/cop-15-l-25-en.pdf

Chávez-Hidalgo, A. (2017). Influencia de la complejidad del hábitat sobre la comunidad bentónica de Bahía Asunción BCS, México Instituto Politécnico Nacional. Centro Interdisciplinario de Ciencias Marinas].

Checkley Jr, D. M., & Barth, J. A. (2009). Patterns and processes in the California Current System. Progress in Oceanography, 83(1-4), 49–64.

Cinner, J. E., Maire, E., Huchery, C., MacNeil, M. A., Graham, N. A., Mora, C., McClanahan, T. R., Barnes, M. L., Kittinger, J. N., & Hicks, C. C. (2018). Gravity of human impacts mediates coral reef conservation gains. Proceedings of the National Academy of Sciences, 115(27), E6116–E6125.

Cooley, S., Schoeman, D., Bopp, L., Boyd, P., Donner, S., Ito, S.-i., Kiessling, W., Martinetto, P., Ojea, E., & Racault, M.-F. (2022). Oceans and Coastal Ecosystems and their Services. In IPCC AR6 WGII. Cambridge University Press.

Donovan, M. K., Burkepile, D. E., Kratochwill, C., Shlesinger, T., Sully, S., Oliver, T. A., Hodgson, G., Freiwald, J., & van Woesik, R. (2021). Local conditions magnify coral loss after marine heatwaves. Science, 372(6545), 977–980.

Dudgeon, S. R., Aronson, R. B., Bruno, J. F., & Precht, W. F. (2010). Phase shifts and stable states on coral reefs. Marine Ecology Progress Series, 413, 201–216.

Edwards, M., & Hernandez-Carmona, G. (2005). Delayed recovery of giant kelp near its southern range limit in the North Pacific following El Niño. Marine Biology, 147, 273–279.

Eger, A. M., Marzinelli, E. M., Christie, H., Fagerli, C. W., Fujita, D., Gonzalez, A. P., Hong, S. W., Kim, J. H., Lee, L. C., & McHugh, T. A. (2022). Global kelp forest restoration: past lessons, present status, and future directions. Biological Reviews, 97(4), 1449–1475.

Filbee-Dexter, K., & Scheibling, R. E. (2014). Sea urchin barrens as alternative stable states of collapsed kelp ecosystems. Marine Ecology Progress Series, 495, 1–25.

Filbee-Dexter, K., Wernberg, T., Grace, S. P., Thormar, J., Fredriksen, S., Narvaez, C., Feehan, C. J., & Norderhaug, K. M. (2020). Marine heatwaves and the collapse of marginal North Atlantic kelp forests. Scientific reports, 10(1), 13388.

Freiwald, J., McMillan, S., & Abbot, D. (2021). Reef check california instruction manual: A guide to monitoring california’s kelp forests. In: Reef Check Foundation, Marina del Rey, CA, USA.

García-Pantoja, J., Olguín-Jacobson, C., Micheli, F., Cavanaugh, K., Montaño-Moctezuma, G., & Arafeh-Dalmau, N. (2025). Manual monitoreo de ecosistemas de bosques de kelp en la Península de Baja California, *México*.

Garrabou, J., Gómez-Gras, D., Medrano, A., Cerrano, C., Ponti, M., Schlegel, R., Bensoussan, N., Turicchia, E., Sini, M., & Gerovasileiou, V. (2022). Marine heatwaves drive recurrent mass mortalities in the Mediterranean Sea. Global Change Biology, 28(19), 5708–5725.

Gove, J. M., Williams, G. J., Lecky, J., Brown, E., Conklin, E., Counsell, C., Davis, G., Donovan, M. K., Falinski, K., & Kramer, L. (2023). Coral reefs benefit from reduced land–sea impacts under ocean warming. Nature, 621(7979), 536–542.

Graham, M. H. (2004). Effects of local deforestation on the diversity and structure of southern California giant kelp forest food webs. Ecosystems, 7, 341–357.

Harden, M., Kovalev, M., Molano, G., Yorke, C., Miller, R., Reed, D., Alberto, F., Koos, D. S., Lansford, R., & Nuzhdin, S. (2024). Heat stress analysis suggests a genetic basis for tolerance in Macrocystis pyrifera across developmental stages. Communications Biology, 7(1), 1147.

Hartig, F. (2018). DHARMa: residual diagnostics for hierarchical (multi-level/mixed) regression models. R Packag version 020.

Hobday, A. J., Alexander, L. V., Perkins, S. E., Smale, D. A., Straub, S. C., Oliver, E. C., Benthuysen, J. A., Burrows, M. T., Donat, M. G., & Feng, M. (2016). A hierarchical approach to defining marine heatwaves. Progress in Oceanography, 141, 227–238.

Hobday, A. J., & Pecl, G. T. (2014). Identification of global marine hotspots: sentinels for change and vanguards for adaptation action. Reviews in Fish Biology and Fisheries, 24(2), 415–425.

Huang, B., Liu, C., Banzon, V., Freeman, E., Graham, G., Hankins, B., Smith, T., & Zhang, H.-M. (2021). Improvements of the daily optimum interpolation sea surface temperature (DOISST) version 2.1. Journal of Climate, 34(8), 2923–2939.

Hughes, B. B., Beas-Luna, R., Barner, A. K., Brewitt, K., Brumbaugh, D. R., Cerny-Chipman, E. B., Close, S. L., Coblentz, K. E., De Nesnera, K. L., & Drobnitch, S. T. (2017). Long-term studies contribute disproportionately to ecology and policy. BioScience, 67(3), 271–281.

IPCC. (2023). Sections. In: Climate Change 2023: Synthesis Report. Contribution of Working Groups I, II and III to the Sixth Assessment Report of the Intergovernmental Panel on Climate Change [Core Writing Team, H. Lee and J. Romero (eds.)].

Jayathilake, D. R., & Costello, M. J. (2021). Version 2 of the world map of laminarian kelp benefits from more Arctic data and makes it the largest marine biome. Biological Conservation, 257, 109099.

Koenker, R. (2023). quantreg: Quantile Regression. R package version 5.95. https://CRAN.R-project.org/package=quantreg.

Kumagai, J. A., Goodman, M. C., Villaseñor-Derbez, J. C., Schoeman, D. S., Cavanuagh, K. C., Bell, T. W., Micheli, F., De Leo, G., & Arafeh-Dalmau, N. (2024). Marine Protected Areas That Preserve Trophic Cascades Promote Resilience of Kelp Forests to Marine Heatwaves. Global Change Biology, 30(12), e17620.

Le, D. M., Desmond, M. J., Pritchard, D. W., & Hepburn, C. D. (2022). Effect of temperature on sporulation and spore development of giant kelp (Macrocystis pyrifera). PLoS One, 17(12), e0278268.

Ling, S., Johnson, C., Frusher, S., & Ridgway, K. (2009). Overfishing reduces resilience of kelp beds to climate-driven catastrophic phase shift. Proceedings of the National Academy of Sciences, 106(52), 22341–22345.

Ling, S., Scheibling, R., Rassweiler, A., Johnson, C., Shears, N., Connell, S., Salomon, A., Norderhaug, K., Pérez-Matus, A., & Hernández, J. (2015). Global regime shift dynamics of catastrophic sea urchin overgrazing. Philosophical Transactions of the Royal Society B: Biological Sciences, 370(1659), 20130269.

Ling, S. D., & Keane, J. P. (2024). Climate-driven invasion and incipient warnings of kelp ecosystem collapse. Nature Communications, 15(1), 400.

Lonhart, S. I., Jeppesen, R., Beas-Luna, R., Crooks, J. A., & Lorda, J. (2019). Shifts in the distribution and abundance of coastal marine species along the eastern Pacific Ocean during marine heatwaves from 2013 to 2018. Marine Biodiversity Records, 12(1), 1–15.

López-de-Lacalle, J. (2016). tsoutliers R package for detection of outliers in time series. *CRAN*, R Package, 95.

Malone, D. P., Davis, K., Lonhart, S. I., Parsons-Field, A., Caselle, J. E., & Carr, M. H. (2022). Large-scale, multidecade monitoring data from kelp forest ecosystems in California and Oregon (USA). In: Wiley Online Library.

McCay, B. J., Micheli, F., Ponce-Díaz, G., Murray, G., Shester, G., Ramirez-Sanchez, S., & Weisman, W. (2014). Cooperatives, concessions, and co-management on the Pacific coast of Mexico. Marine Policy, 44, 49–59.

McPherson, M. L., Finger, D. J. I., Houskeeper, H. F., Bell, T. W., Carr, M. H., Rogers-Bennett, L., & Kudela, R. M. (2021). Large-scale shift in the structure of a kelp forest ecosystem co-occurs with an epizootic and marine heatwave. Communications Biology, 4(1), 298. 10.1038/s42003-021-01827-6

Micheli, F., Saenz-Arroyo, A., Aalto, E., Beas-Luna, R., Boch, C. A., Cardenas, J. C., De Leo, G. A., Diaz, E., Espinoza-Montes, A., Finkbeiner, E., Freiwald, J., Fulton, S., Hernández, A., Lejbowicz, A., Low, N. H. N., Martinez, R., McCay, B., Monismith, S., Precoma-de la Mora, M., . . . Woodson, C. B. (2024). Social-ecological vulnerability to environmental extremes and adaptation pathways in small-scale fisheries of the southern California Current [Original Research]. Frontiers in Marine Science, 11. 10.3389/fmars.2024.1322108

Micheli, F., Saenz-Arroyo, A., Greenley, A., Vazquez, L., Espinoza Montes, J. A., Rossetto, M., & De Leo, G. A. (2012). Evidence that marine reserves enhance resilience to climatic impacts. PloS one, 7(7), e40832.

Olguín-Jacobson, C., Arafeh-Dalmau, N., Early-Capistrán, M.-M., Kumagai, J. A., Schoeman, D. S., Espinoza Montes, J. A., Hernández-Velasco, A., Martínez, R., Romero, A., Torre, J., Woodson, C. B., De Leo, G., & Micheli, F. (2025). Recovery mode: Marine protected areas enhance climate resilience of invertebrate species to marine heatwaves. Functional Ecology, n/a(n/a). 10.1111/1365-2435.70060

Oliver, E. C., Burrows, M. T., Donat, M. G., Sen Gupta, A., Alexander, L. V., Perkins-Kirkpatrick, S. E., Benthuysen, J. A., Hobday, A. J., Holbrook, N. J., & Moore, P. J. (2019). Projected marine heatwaves in the 21st century and the potential for ecological impact. Frontiers in Marine Science, 6, 734.

Oliver, E. C. J., Donat, M. G., Burrows, M. T., Moore, P. J., Smale, D. A., Alexander, L. V., Benthuysen, J. A., Feng, M., Sen Gupta, A., Hobday, A. J., Holbrook, N. J., Perkins-Kirkpatrick, S. E., Scannell, H. A., Straub, S. C., & Wernberg, T. (2018). Longer and more frequent marine heatwaves over the past century [Article]. Nature Communications, 9(1), Article 1324. 10.1038/s41467-018-03732-9

Oliver, T. H., Heard, M. S., Isaac, N. J., Roy, D. B., Procter, D., Eigenbrod, F., Freckleton, R., Hector, A., Orme, C. D. L., & Petchey, O. L. (2015). Biodiversity and resilience of ecosystem functions. Trends in ecology & evolution, 30(11), 673–684.

Pecl, G. T., Araújo, M. B., Bell, J. D., Blanchard, J., Bonebrake, T. C., Chen, I.-C., Clark, T. D., Colwell, R. K., Danielsen, F., & Evengård, B. (2017). Biodiversity redistribution under climate change: Impacts on ecosystems and human well-being. Science, 355(6332), eaai9214.

Pinsky, M. L., Selden, R. L., & Kitchel, Z. J. (2020). Climate-driven shifts in marine species ranges: scaling from organisms to communities. Annual Review of Marine Science, 12(1), 153–179.

Poloczanska, E. S., Brown, C. J., Sydeman, W. J., Kiessling, W., Schoeman, D. S., Moore, P. J., Brander, K., Bruno, J. F., Buckley, L. B., & Burrows, M. T. (2013). Global imprint of climate change on marine life. Nature Climate Change, 3(10), 919–925.

Poloczanska, E. S., Burrows, M. T., Brown, C. J., García Molinos, J., Halpern, B. S., Hoegh-Guldberg, O., Kappel, C. V., Moore, P. J., Richardson, A. J., Schoeman, D. S., & Sydeman, W. J. (2016). Responses of Marine Organisms to Climate Change across Oceans [Review]. Frontiers in Marine Science, 3(62). 10.3389/fmars.2016.00062

Ramirez-Valdez, A., Aburto-Oropeza, O., Palacios-Salgado, D. S., Correa-Sandoval, F., Ramirez-Valdez, A., Carlos Villasenor-Derbez, J., Jose Cota-Nieto, J., Hinojosa-Arango, G., Reyes-Bonilla, H., & Dominguez-Guerrero, I. (2015). The nearshore fishes of the Cedros archipelago (North-Eastern Pacific) and their biogeographic affinities. California Cooperative Oceanic Fisheries Investigations Reports, 56, 143–167.

Rogers-Bennett, L., & Catton, C. A. (2019). Marine heat wave and multiple stressors tip bull kelp forest to sea urchin barrens. Scientific reports, 9(1), 15050. 10.1038/s41598-019-51114-y

Rothäusler, E., Gomez, I., Hinojosa, I. A., Karsten, U., Tala, F., & Thiel, M. (2011). PHYSIOLOGICAL PERFORMANCE OF FLOATING GIANT KELP MACROCYSTIS PYRIFERA (PHAEOPHYCEAE): LATITUDINAL VARIABILITY IN THE EFFECTS OF TEMPERATURE AND GRAZING 1. Journal of Phycology, 47(2), 269–281.

Sanford, E., Sones, J. L., García-Reyes, M., Goddard, J. H. R., & Largier, J. L. (2019). Widespread shifts in the coastal biota of northern California during the 2014–2016 marine heatwaves. Scientific reports, 9(1), 4216. 10.1038/s41598-019-40784-3

Schlegel, R. W., & Smit, A. J. (2018). heatwaveR: A central algorithm for the detection of heatwaves and cold-spells. Journal of Open Source Software, 3(27), 821.

Schmeller, D. S., Weatherdon, L. V., Loyau, A., Bondeau, A., Brotons, L., Brummitt, N., Geijzendorffer, I. R., Haase, P., Kuemmerlen, M., & Martin, C. S. (2018). A suite of essential biodiversity variables for detecting critical biodiversity change. Biological Reviews, 93(1), 55–71.

Smale, D. A. (2020). Impacts of ocean warming on kelp forest ecosystems. New Phytologist, 225(4), 1447–1454.

Smale, D. A., Wernberg, T., Oliver, E. C. J., Thomsen, M., Harvey, B. P., Straub, S. C., Burrows, M. T., Alexander, L. V., Benthuysen, J. A., Donat, M. G., Feng, M., Hobday, A. J., Holbrook, N. J., Perkins-Kirkpatrick, S. E., Scannell, H. A., Sen Gupta, A., Payne, B. L., & Moore, P. J. (2019). Marine heatwaves threaten global biodiversity and the provision of ecosystem services. Nature Climate Change, 9(4), 306–312. 10.1038/s41558-019-0412-1

Smith, J. G., Malone, D., & Carr, M. H. (2024). Consequences of kelp forest ecosystem shifts and predictors of persistence through multiple stressors. Proceedings of the Royal Society B: Biological Sciences, 291(2016), 20232749. doi:10.1098/rspb.2023.2749

Smith, K. E., Aubin, M., Burrows, M. T., Filbee-Dexter, K., Hobday, A. J., Holbrook, N. J., King, N. G., Moore, P. J., Sen Gupta, A., & Thomsen, M. (2024). Global impacts of marine heatwaves on coastal foundation species. Nature Communications, 15(1), 5052.

Smith, K. E., Burrows, M. T., Hobday, A. J., King, N. G., Moore, P. J., Sen Gupta, A., Thomsen, M. S., Wernberg, T., & Smale, D. A. (2023). Biological impacts of marine heatwaves. Annual Review of Marine Science, 15, 119–145.

Smith, K. E., Burrows, M. T., Hobday, A. J., Sen Gupta, A., Moore, P. J., Thomsen, M., Wernberg, T., & Smale, D. A. (2021). Socioeconomic impacts of marine heatwaves: Global issues and opportunities. Science, 374(6566), eabj3593.

Starko, S., Epstein, G., Chalifour, L., Bruce, K., Buzzoni, D., Csordas, M., Dimoff, S., Hansen, R., Maucieri, D., & McHenry, J. (2025). Ecological responses to extreme climatic events: a systematic review of the 2014-2016 Northeast Pacific marine heatwave. Oceanogr. Mar. Biol. Annu. Rev.

Torda, G., Donelson, J. M., Aranda, M., Barshis, D. J., Bay, L., Berumen, M. L., Bourne, D. G., Cantin, N., Foret, S., & Matz, M. (2017). Rapid adaptive responses to climate change in corals. Nature Climate Change, 7(9), 627–636.

Vergés, A., Steinberg, P. D., Hay, M. E., Poore, A. G., Campbell, A. H., Ballesteros, E., Heck Jr, K. L., Booth, D. J., Coleman, M. A., & Feary, D. A. (2014). The tropicalization of temperate marine ecosystems: climate-mediated changes in herbivory and community phase shifts. Proceedings of the Royal Society B: Biological Sciences, 281(1789), 20140846.

Villaseñor-Derbez, J. C., Arafeh-Dalmau, N., & Micheli, F. (2024). Past and future impacts of marine heatwaves on small-scale fisheries in Baja California, Mexico. Communications Earth & Environment, 5(1), 623.

Villaseñor-Derbez, J. C., Arafeh-Dalmau, N., Micheli, F. (2024). Imapcts of past and future marine heatwaves in Baja California, Mexico [Article]. Communications Earth & Environment. 10.1038/s43247-024-01696-x

Vranken, S., Wernberg, T., Scheben, A., Pessarrodona, A., Batley, J., & Coleman, M. A. (2025). Connectivity enhances resilience of marine forests after an extreme event. Scientific reports, 15(1), 5019.

Webster, M. M., Twohey, B., Alagona, P. S., Arafeh-Dalmau, N., Colton, M. A., De Stigter, E., Eger, A. M., Miller, S. N., Pecl, G. T., & Scheffers, B. R. (2023). Assisting adaptation in a changing world. Frontiers in Environmental Science, 11, 1232374.

Wernberg, T. (2021). Marine heatwave drives collapse of kelp forests in Western Australia. In Ecosystem collapse and climate change (pp. 325-343). Springer.

Wernberg, T., Bennett, S., Babcock, R. C., De Bettignies, T., Cure, K., Depczynski, M., Dufois, F., Fromont, J., Fulton, C. J., & Hovey, R. K. (2016). Climate-driven regime shift of a temperate marine ecosystem. Science, 353(6295), 169–172.

Wernberg, T., Coleman, M. A., Babcock, R. C., Bell, S. Y., Bolton, J. J., Connell, S. D., Hurd, C. L., Johnson, C. R., Marzinelli, E. M., & Shears, N. T. (2019). Biology and ecology of the globally significant kelp Ecklonia radiata. Oceanography and marine biology.

Wernberg, T., Coleman, M. A., Bennett, S., Thomsen, M. S., Tuya, F., & Kelaher, B. P. (2018). Genetic diversity and kelp forest vulnerability to climatic stress. Scientific reports, 8(1), 1851.

Wernberg, T., Thomsen, M. S., Burrows, M. T., Filbee-Dexter, K., Hobday, A. J., Holbrook, N. J., Montie, S., Moore, P. J., Oliver, E. C. J., Sen Gupta, A., Smale, D. A., & Smith, K. (2025). Marine heatwaves as hot spots of climate change and impacts on biodiversity and ecosystem services. Nature Reviews Biodiversity, 1(7), 461–479. 10.1038/s44358-025-00058-5

White, J. W., Hopf, J. K., Arafeh-Dalmau, N., Ban, N. C., Bates, A. E., Claudet, J., Lopazanski, C., Sunday, J. M., & Caselle, J. E. (2025). Measurements, mechanisms, and management recommendations for how marine protected areas can provide climate resilience. Marine Policy, 171, 106419.

Woodson, C. B., Micheli, F., Boch, C., Al-Najjar, M., Espinoza, A., Hernandez, A., Vázquez-Vera, L., Saenz-Arroyo, A., Monismith, S. G., & Torre, J. (2019). Harnessing marine microclimates for climate change adaptation and marine conservation. Conservation Letters, 12(2), e12609.

