## Supplementary Figures and Tables for "Marine heatwaves drive range contraction and alternative states of kelp forests at their warm limit"

### Supplementary material

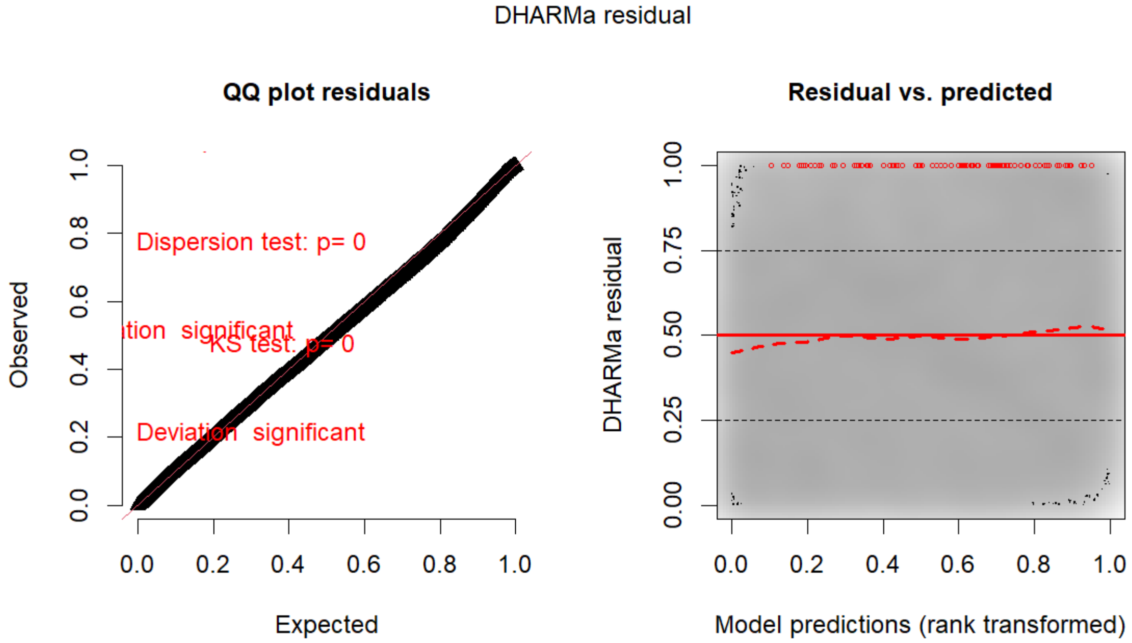


**Figure S1:** Residuals plots from the DHARMa package form a Tweedie GAMM that predicts remote-sensed kelp canopy area based on the interaction of ecoregion (northern and central Baja California) and year (as a smooth term), including a spatial random field. (Figure 2a in the main text).

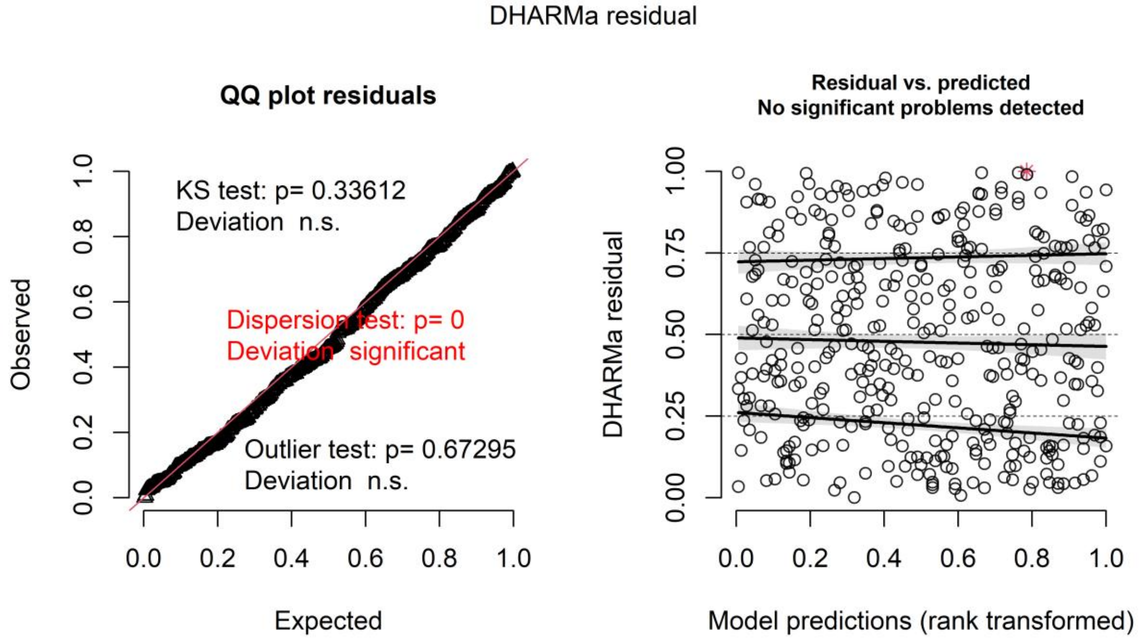


**Figure S2:** Residuals plots from the DHARMa package form a Nbinom1 GLMM that predicts giant kelp abundance based on the interaction of ecoregion (northern or central Baja California) and year, including a spatial random field.

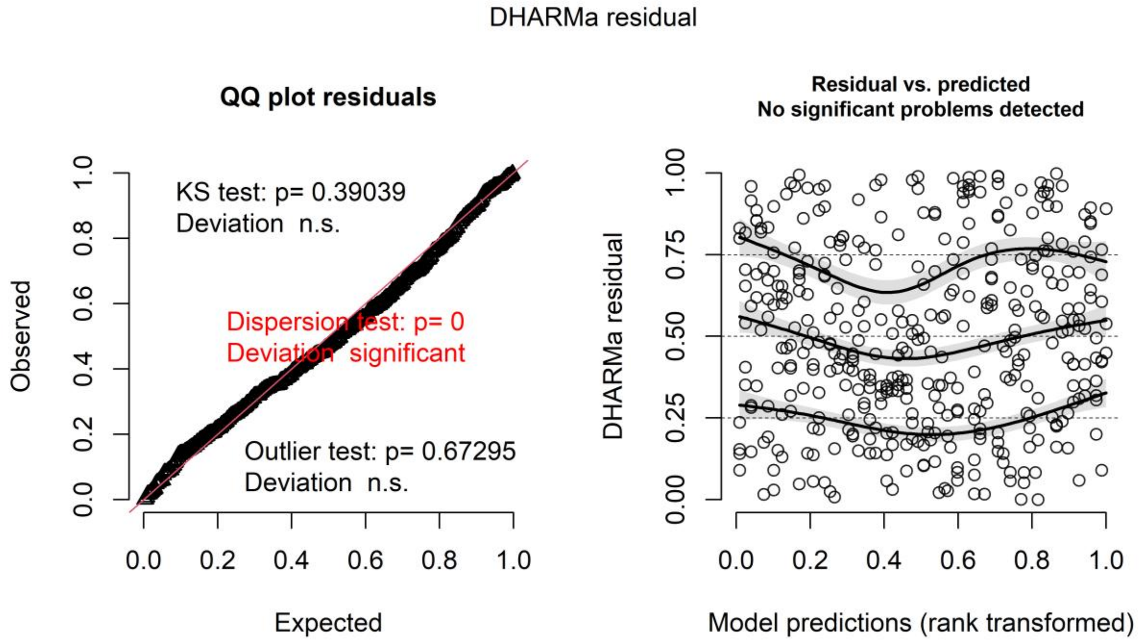


**Figure S3:** Residuals plots from the DHARMa package form a Nbinom1 GLMM that predicts palm kelp abundance based on the ecoregion (northern or central Baja California), including a spatial random field.

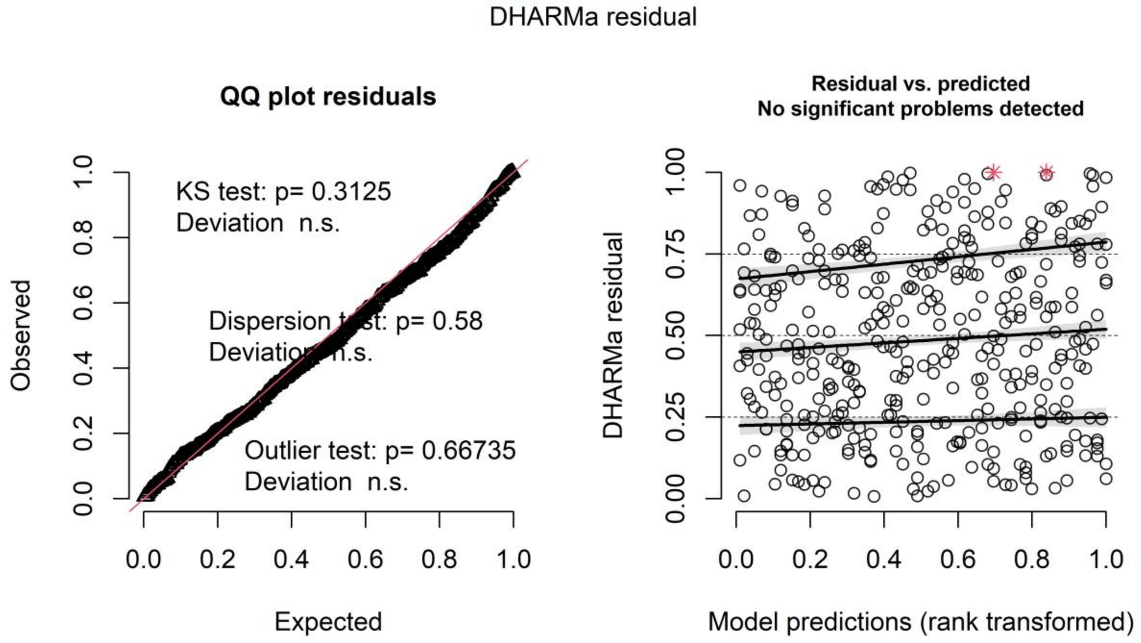


**Figure S4:** Residuals plots from the DHARMa package form a Nbinom1 GLMM that predicts lobster abundance based on ecoregion (northern or central Baja California), including a spatial random field.

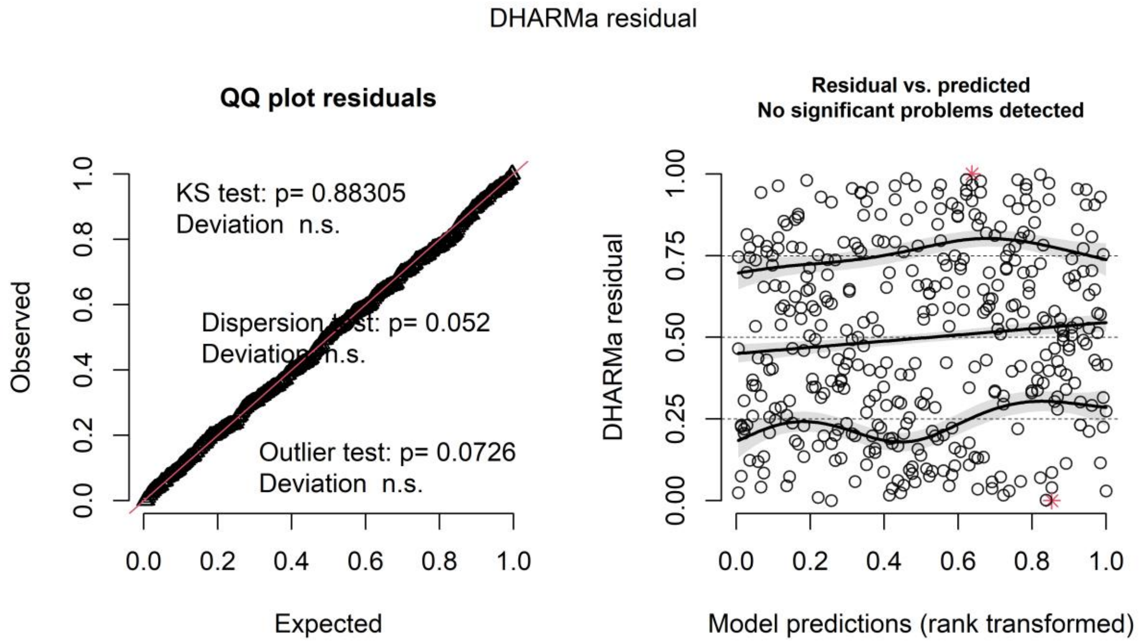


**Figure S5:** Residuals plots from the DHARMa package form a Nbinom1 GLMM that predicts large (> 35cm length) sheepshead biomass based on the interaction of ecoregion (northern or central Baja California) and year, including a spatial random field.

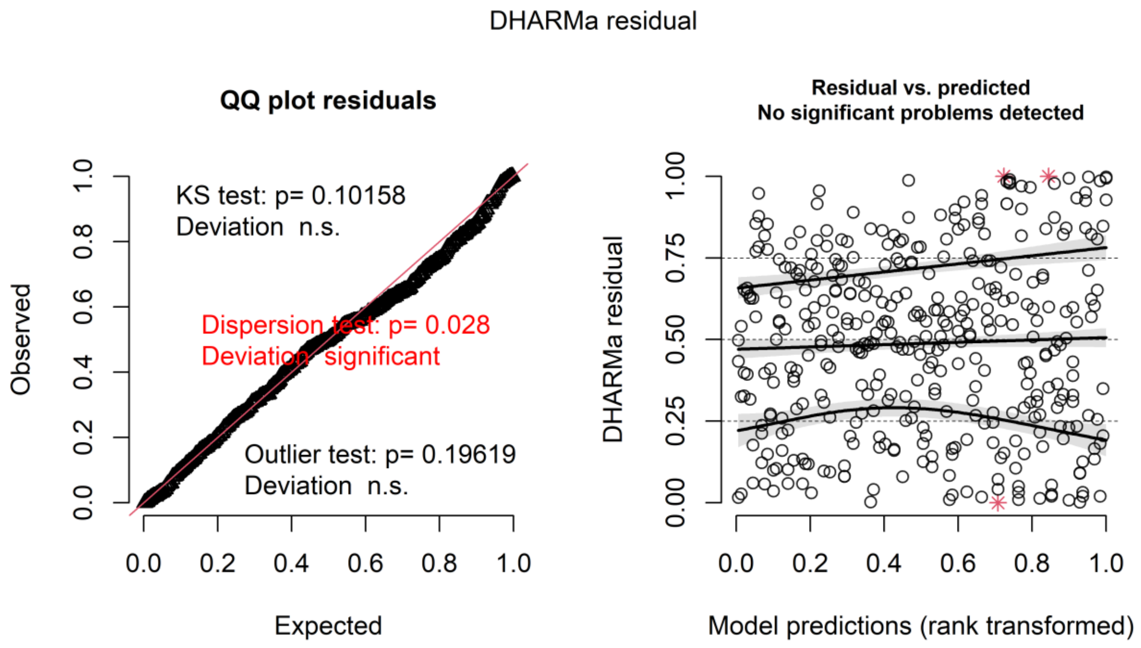


**Figure S6:** Residuals plots from the DHARMa package form a Tweedie GLMM that predicts urchin abundance based on the interaction of ecoregion (northern or central Baja California) and year, including a spatial random field.

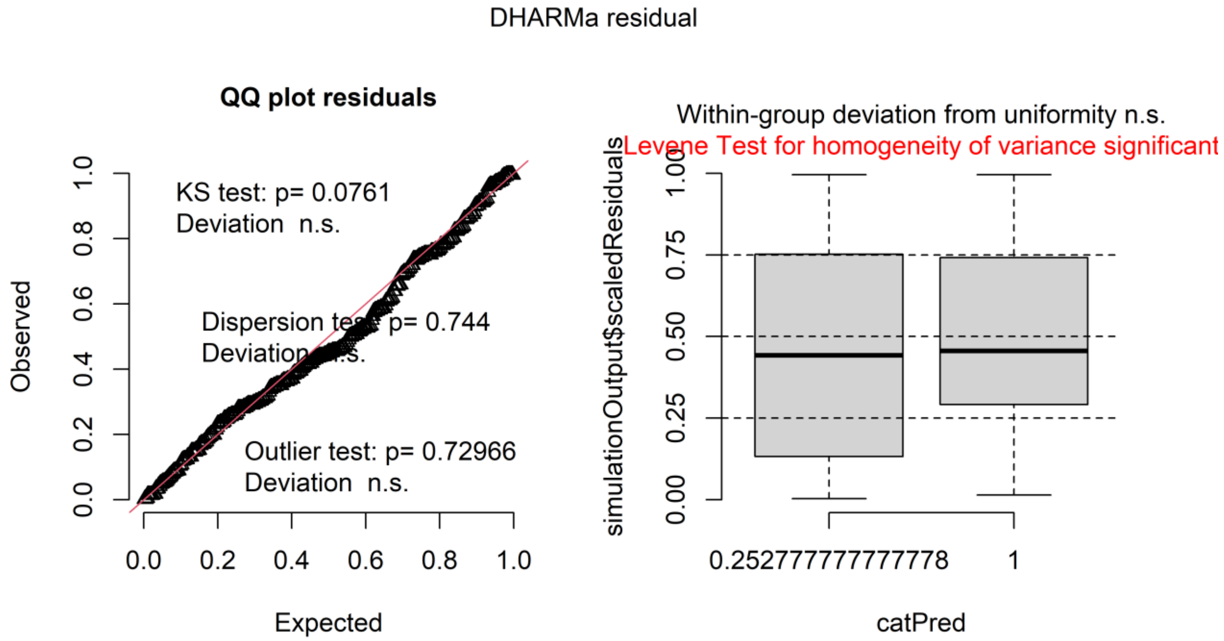


**Figure S7:** Residuals plots from the DHARMa package form a Tweedie GLMM that predicts urchin density based on marine heatwave period (before vs after) for the Ensenada region.

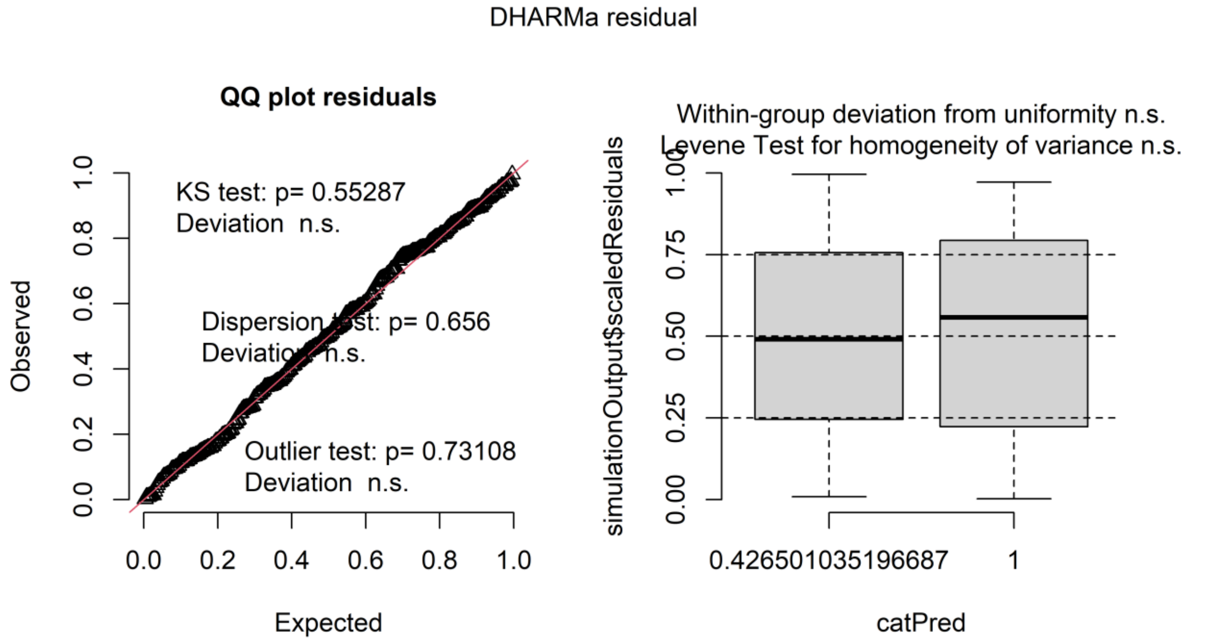


.

**Figure S8:** Residuals plots from the DHARMa package form a Tweedie GLMM that predicts palm kelp density based on marine heatwave period (before vs after) for the Ensenada region

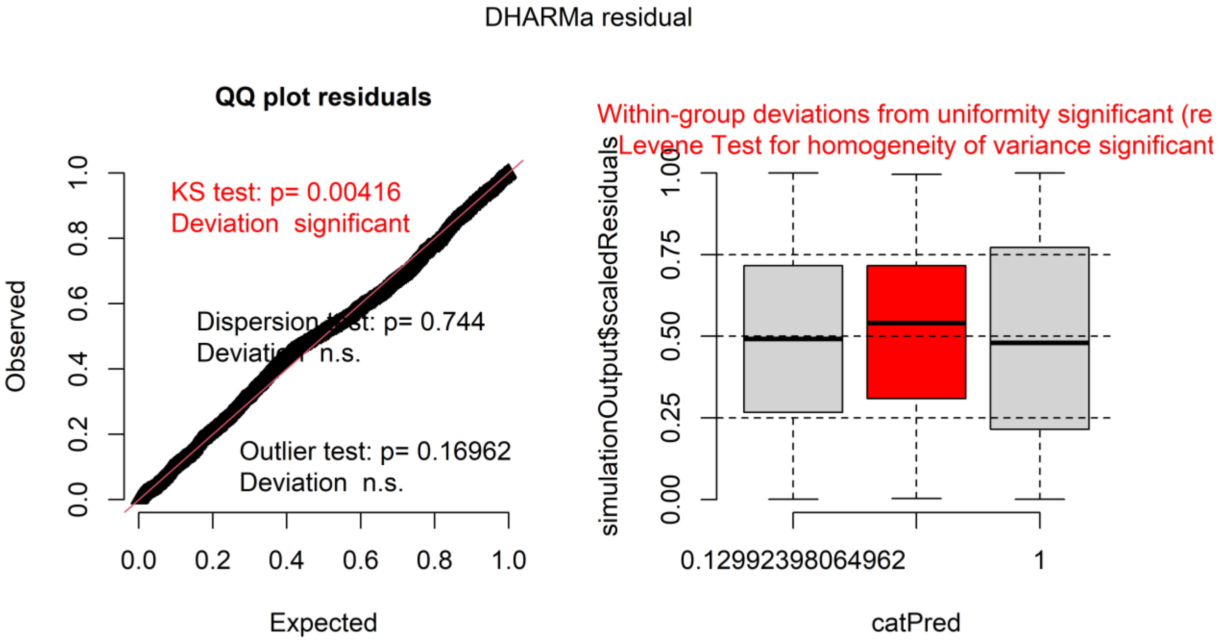


**Figure S9:** Residuals plots from the DHARMa package form a Tweedie GLMM that predicts urchin density based on marine heatwave period (before, during, and after) for the Eugenia region, including temporal autocorrelation (AR1).

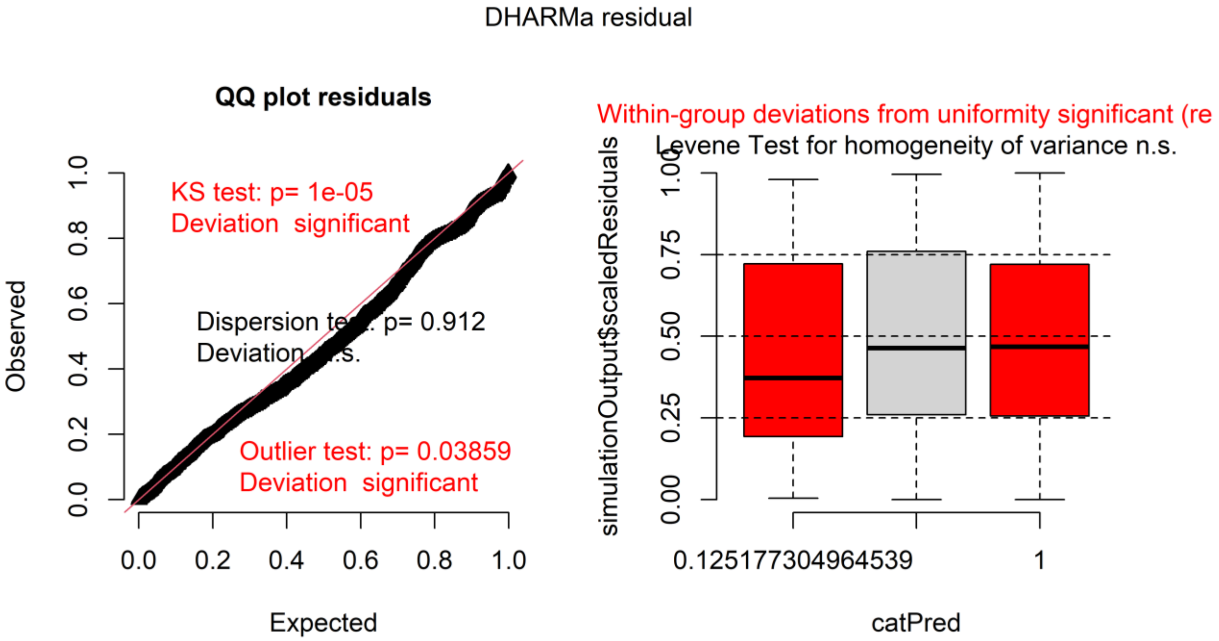


**Figure S10**: Residuals plots from the DHARMa package form a Tweedie GLMM that predicts palm kelp density based on marine heatwave period (before, during, and after) for the Eugenia region, including temporal autocorrelation (AR1).

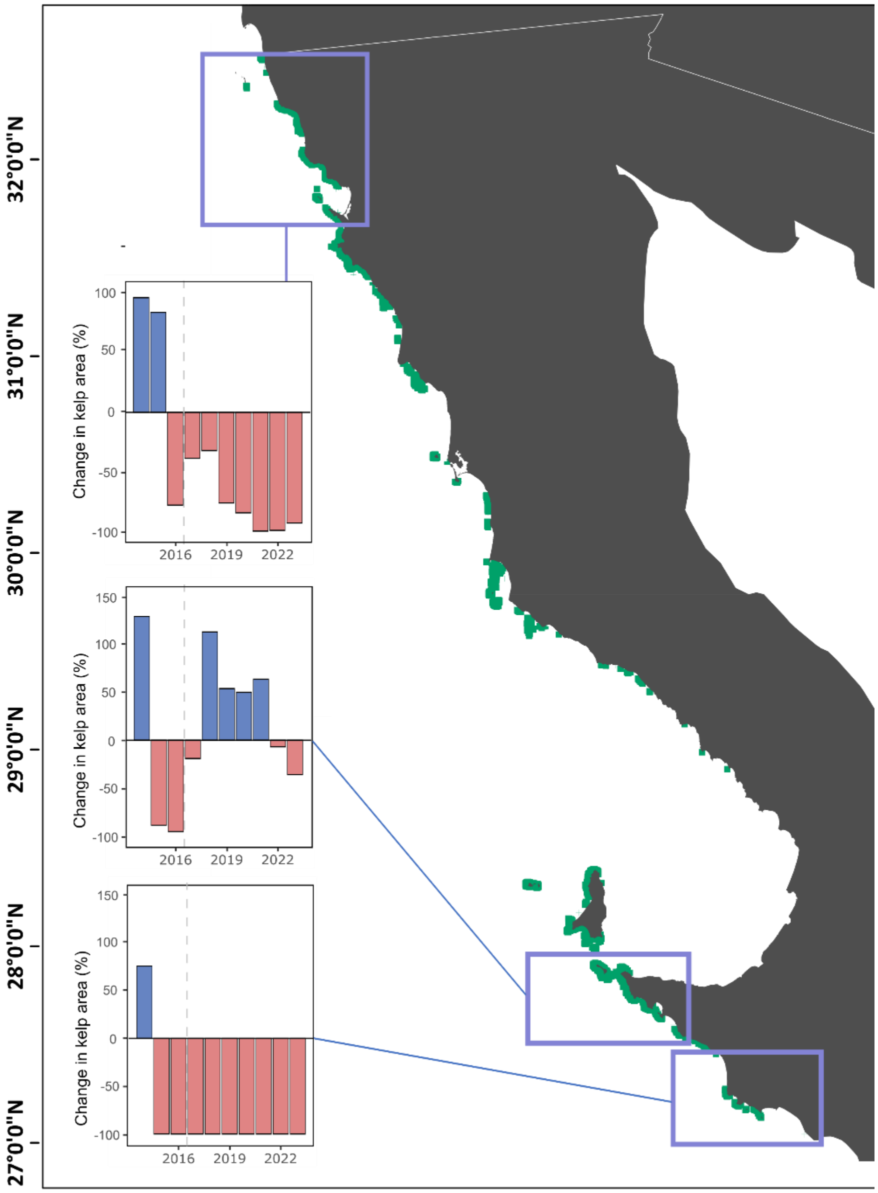


**Figure S11.** Kelp forests in Baja California, Mexico, and their dynamics following the 2014–2016 marine heatwaves (MHWs). Bar plots show annual percentage change in kelp cover relative to the 2000–2013 baseline, binned (0.5°) by region. Blue bars indicate years above the baseline, and red bars indicate years below.

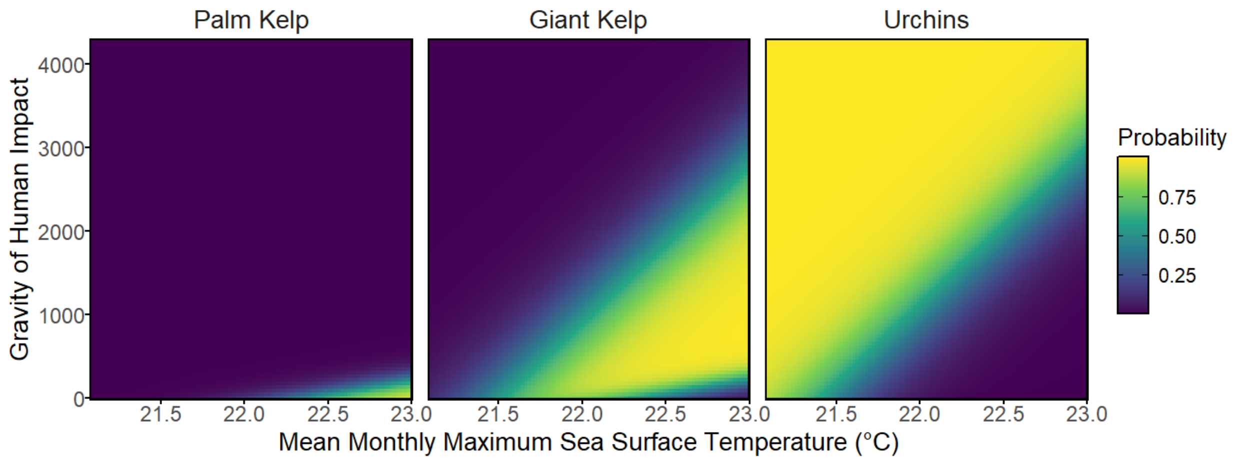


**Figure S2.** Predicted probabilities of alternative ecological states across gradients of temperature and human impact. Each panel shows the modeled probability of dominance for palm kelp, giant kelp, and urchins across 72 sub-site observations (36 sites with two sub-sites) aggregated for 2022–2023 in Baja California, Mexico. Probabilities are derived from a multinomial model incorporating environmental predictors, with outputs visualized across the environmental space. Mean monthly maximum sea surface temperature (°C) (after MHWs: 2017-2023) and gravity of human impact are continuous variables, while color gradients indicate the relative probability of each state, ranging from low (purple) to high (yellow).

**Table S1.** Surveyed sites included in this manuscript.

| Site | Region | Program | Years |
|---|---|---|---|
| Bajo Coronado | Northern Baja California | UABC/MasKelp | 2011-2013, 2019-2023 |
| Campo Lopez | Northern Baja California | UABC/MasKelp | 2011-2013, 2019-2023 |
| Baja Mar | Northern Baja California | UABC/MasKelp | 2011-2013, 2019-2023 |
| Sal Si Puedes | Northern Baja California | UABC/MasKelp | 2011-2013, 2019-2023 |
| San Miguel | Northern Baja California | UABC/MasKelp | 2011-2013, 2019-2023 |
| Isla Todos Santos Expuesto | Northern Baja California | UABC/MasKelp | 2011-2013, 2016, 2019-2023 |
| Isla Todos Santos Protegido | Northern Baja California | UABC/MasKelp | 2011-2013, 2016, 2019-2023 |
| Punta Banda | Northern Baja California | MasKelp | 2022-2023 |
| El Retiro | Northern Baja California | UABC/MasKelp | 2011-2013, 2019-2023 |
| La Bocana | Northern Baja California | MasKelp | 2022-2023 |
| Erendira | Northern Baja California | UABC/MasKelp | 2011-2013, 2019-2022 |
| Punta Colonet | Northern Baja California | UABC/MasKelp | 2022 |
| Isla San Maritn Expuesto | Northern Baja California | UABC/MasKelp | 2011-2013, 2016, 2019-2023 |
| Isla San Martin Protegido | Northern Baja California | UABC/MasKelp | 2011-2013, 2016, 2019-2023 |
| El Rosario Picacho | Northern Baja California | MasKelp | 2022-2023 |
| El Rosario Lazaro | Northern Baja California | UABC/MasKelp | 2011-2013, 2019-2023 |
| Isla San Jeronimo Expuesto | Northern Baja California | UABC/MasKelp | 2011-2013, 2016, 2019-2023 |
| Isla San Jeronimo Protegido | Northern Baja California | UABC/MasKelp | 2011-2013, 2016, 2019-2023 |
| Arrecife Sacramento | Northern Baja California | UABC/MasKelp | 2011-2013, 2016, 2019-2021, 2023 |
| Isla Cedros Punta Norte | Central Baja California | UABC/MasKelp | 2019, 2022-2023 |
| Isla Cedros Puerto Escondido | Central Baja California | MasKelp | 2022-2023 |
| Isla Cedros San Agustin | Central Baja California | UABC/MasKelp | 2019, 2022-2023 |
| Isla Cedros La Botella | Central Baja California | MasKelp | 2022-2023 |
| Isla Benitos el Tecapan | Central Baja California | UABC/MasKelp | 2019, 2022-2023 |
| Isla San Benitos Los Huevos | Central Baja California | MasKelp | 2022-2023 |
| Isla San Benitos | Central Baja California | MasKelp | 2022-2023 |
| Isla Natividad La Vela | Central Baja California | MasKelp | 2022-2023 |
| Isla Natividad El Tibo | Central Baja California | MasKelp | 2022-2023 |
| Isla Natividad La Plana | Central Baja California | COBI/Stanford | 2006-2023 |
| Isla Natividad La Guanera | Central Baja California | COBI/Stanford | 2006-2023 |
| Isla Natividad Babencho | Central Baja California | COBI/Stanford | 2006-2023 |
| Isla Natividad | Central Baja California | COBI/Stanford | 2006-2023 |
| Isla Natividad | Central Baja California | COBI/Stanford | 2006-2023 |

| | | | |
|---|---|---|---|
| Punta Eugenia El Reventon | Central Baja California | MasKelp | 2022-2023 |
| Punta Eugenia Las Boyas | Central Baja California | MasKelp | 2022-2023 |
| Bahia Tortugas Clambei | Central Baja California | MasKelp | 2022-2023 |
| Bahia Tortugas La Rompiente | Central Baja California | MasKelp | 2022-2023 |
| Bahia Tortugas | Central Baja California | MasKelp | 2022-2023 |
| Emancipacion Puerto Escondido | Central Baja California | MasKelp | 2022-2023 |
| Emancipacion | Central Baja California | MasKelp | 2022-2023 |
| Punt San Roque | Central Baja California | MasKelp | 2019, 2022-2023 |
| Isla Asunion | Central Baja California | MasKelp | 2019, 2022-2023 |
| San Pablo Puerto | Central Baja California | INAPESCA | 1993-2013 |
| San Pablo Limite | Central Baja California | INAPESCA | 1993-2013 |
| Vuelta del Cerro | Central Baja California | INAPESCA | 1993-2013 |
| El Reef | Central Baja California | INAPESCA | 1993-2013 |
| La Puntilla | Central Baja California | INAPESCA | 1993-2013 |
| El Vapor | Central Baja California | INAPESCA | 1993-2013 |

**Table S2.** Mean cumulative annual marine heatwave intensities (C° days) (± 95% confidence intervals) from 2000-2023 for northern (n = 19) and central Baja California (n= 18) giant kelp forests, based on OISST historical data.

| Year | Northern Baja California | Central Baja California |
|---|---|---|
| 2000 | 18.36 ± 8.10 | 18.84 ± 2.95 |
| 2001 | 24.53 ± 3.61 | 34.14 ± 4.53 |
| 2002 | 0.00 | 0.61 ± 1.19 |
| 2003 | 26.05 ± 6.34 | 10.89 ± 3.01 |
| 2004 | 10.85 ± 4.46 | 8.49 ± 4.93 |
| 2005 | 7.76 ± 3.94 | 7.92 ± 4.32 |
| 2006 | 120.53 ± 15.22 | 96.85 ± 18.23 |
| 2007 | 8.70 ± 4.81 | 3.96 ± 2.37 |
| 2008 | 13.16 ± 8.29 | 76.63 ± 22.00 |
| 2009 | 7.18 ± 2.76 | 4.48 ± 4.50 |
| 2010 | 11.23 ± 6.06 | 26.47 ± 6.49 |
| 2011 | 0.00 | 0.00 |
| 2012 | 27.10 ± 6.94 | 188.13 ± 27.21 |
| 2013 | 34.85 ± 7.36 | 26.57 ± 2.98 |
| 2014 | 427.37 ± 25.86 | 561.23 ± 46.33 |

| | | |
|---|---|---|
| 2015 | 741.64 ± 20.76 | 895.59 ± 21.61 |
| 2016 | 233.10 ± 6.36 | 300.63 ± 27.33 |
| 2017 | 181.86 ± 12.45 | 124.84 ± 21.14 |
| 2018 | 273.12 ± 12.93 | 156.10 ± 28.65 |
| 2019 | 34.06 ± 8.37 | 20.14 ± 7.99 |
| 2020 | 332.93 ± 32.76 | 147.28 ± 28.58 |
| 2021 | 157.46 ± 45.58 | 73.45 ± 16.91 |
| 2022 | 61.48 ± 12.20 | 40.11 ± 11.30 |
| 2023 | 38.23 ± 4.10 | 135.10 ± 14.79 |

**Table S3.-** Estimated means effects for northern and central Baja California regions, Mexico, for 2022-2023 for giant kelp, palm kelp, urchins, lobster, and sheepshead. Values are (individuals/$m^2$ for all species, and biomass for large (size > 35 cm total length) sheepsheads (kg/$m^2$)) for GLMM with spatial autocorrelation.

| Region | Year | Giant kelp response | SE | Palm kelp response | SE | Urchins response | SE | Lobster response | SE | Sheepheads | SE |
|---|---|---|---|---|---|---|---|---|---|---|---|
| Northern | 2022 | 0.1863 | 0.2011 | 0.267 | 0.204 | 71.27 | 23.21 | 0.188 | 0.0654 | 3.32 | 1.44 |
| Central | 2022 | 4.4937 | 3.5587 | 12.136 | 6.711 | 5.37 | 1.65 | 3.431 | 0.6068 | 37.64 | 9.55 |
| Northern | 2023 | 0.0765 | 0.0841 | 0.148 | 0.118 | 190.21 | 61.52 | 0.188 | 0.0654 | 5.27 | 2.20 |
| Central | 2023 | 4.1005 | 3.2584 | 12.559 | 6.965 | 4.65 | 1.49 | 3.431 | 0.6068 | 123.73 | 29.48 |

**Table S4.** Pairwise tests for significant differences between regions (northern and central Baja California, Mexico) for giant kelp, palm kelp, urchins, lobster, and sheepshead. Ratio for GLMM with spatial autocorrelation indicates relative density (individuals/$m^2$ for all species, and biomass for large (size > 35 cm total length) sheepsheads (kg/$m^2$)) across the respective comparisons. A value in the ratio of <1 means an increase in density or biomass, while a value >1 means a decrease in the respective comparison.

| Comparison | Year | Giant kelp | | Palm Kelp | | Lobster | | Sheepheads | | Urchins | |
|---|---|---|---|---|---|---|---|---|---|---|---|
| | | Ratio | p value | Ratio | p value | Ratio | p value | Ratio | p value | Ratio | p value |
| Central/ Northern | 2022 | 24.1 | 0.0122 | 45.5 | <0.0001 | 18.3 | <0.0001 | 11.3 | <0.0001 | 0.0753 | <0.0001 |
| Central/ Northern | 2023 | 53.6 | 0.0020 | 84.9 | <0.0001 | 18.3 | <0.0001 | 23.5 | <0.0001 | 0.0244 | <0.0001 |

**Table S5**. ARIMA outliers for the Ensenada region with time and type of outlier from 1991-2023. Outlier types are Additive outlier (AO), Temporal change (TC), and Level Shift (LS).

| Number | Type | Year | Quarter | Coefhat | tstat |
|---|---|---|---|---|---|
| 1 | TC | 1994 | 2 | 974609.7 | 4.374 |
| 2 | TC | 1994 | 4 | -788045.5 | -3.411 |
| 3 | AO | 2001 | 2 | 1211261.6 | 6.843 |
| 4 | AO | 2007 | 2 | 914006.8 | 5.794 |
| 5 | TC | 2008 | 2 | 708053.3 | 3.476 |
| 6 | TC | 2009 | 1 | 882324.9 | 4.25 |
| 7 | AO | 2009 | 2 | 1429721.6 | 8.409 |
| 8 | AO | 2010 | 3 | 1086017.2 | 6.340 |
| 9 | TC | 2011 | 2 | 874162.9 | 4.261 |
| 10 | TC | 2013 | 2 | 1081501.6 | 5.220 |
| 11 | TC | 2014 | 2 | 914533.6 | 4.303 |
| 12 | AO | 2014 | 3 | 961781.1 | 5.606 |
| 13 | LS | 2015 | 2 | 799184.1 | 3.407 |
| 14 | AO | 2015 | 3 | 965713.2 | 5.426 |
| 15 | LS | 2015 | 4 | -1054793.6 | -4.327 |
| 16 | TC | 2017 | 3 | 871229.4 | 4.255 |

**Table S6.** ARIMA outliers for the Punta Eugenia region with time and type of outlier from 2000-2023. Outlier types are Additive outlier (AO), Temporal change (TC), and Level Shift (LS).

| Number | Type | Year | Quarter | Coefhat | tstat |
|---|---|---|---|---|---|
| 1 | AO | 2014.25 | AO | 7044536 | 3.625 |
| 2 | AO | 2018.25 | AO | 10006139 | 5.413 |

**Table S7**. ARIMA outliers for the Bahia Asuncion region with time and type of outlier from 1993-2023. Outlier types are Additive outlier (AO), Temporal change (TC), and Level Shift (LS).

| Number | Type | Year | Quarter | Coefhat | tstat |
|---|---|---|---|---|---|
| 1 | LS | 2001.50 | 3 | 329776.1 | 3.630 |
| 2 | AO | 2008.25 | 2 | 1313419.5 | 6.2227 |
| 3 | LS | 2011.00 | 1 | 813731.8 | 3.842 |
| 4 | LS | 2011.50 | 3 | -847236.3 | -3.907 |
| 5 | LS | 2014.75 | 4 | -438576.3 | -3.708 |
